# Metabolomic analysis of the effect of endocannabinoid metabolism inhibition in ovalbumin-induced allergic airway inflammation in Guinea pigs

**DOI:** 10.1101/2024.03.10.584309

**Authors:** Reshed Abohalaka, Yasemin Karaman, Tuba Recber, Sevgen Celik Onder, Emirhan Nemutlu, Turgut Emrah Bozkurt

## Abstract

**Background and aims:** Asthma manifests as a multifaceted airway inflammation. The therapeutic potential of targeting endocannabinoids in mitigating asthma remains incompletely elucidated. Therefore, we aim to scrutinize metabolic alterations, deepen our comprehension of the endocannabinoids’ therapeutic role, and discern novel biomarkers for monitoring allergic airway inflammation.

**Methods:** Guinea pigs were sensitized with ovalbumin (150μg) or PBS on days 1, 4, and 7. On day 14, they were exposed to aerosols containing 0.3% ovalbumin or PBS. Treatment groups were administered inhibitors of fatty acid amide hydrolase (FAAH) and/or monoacylglycerol lipase (MAGL) one hour prior to aerosol exposure. Subsequently, bronchoalveolar lavage (BAL), blood, and lung samples were collected on the following day for analysis using GC-MS. Metabolites were meticulously categorized into 10 distinct classifications based on their chemical and biological functions. Subsequently, they were further organized into 5 principal metabolic pathways for a comprehensive metabolomic analysis.

**Results:** Ovalbumin exposure exclusively altered the metabolic profile in the lung. Conversely, inhibition of endocannabinoids metabolism induced a systemic shift in energy metabolites such as carbohydrates, amino and fatty acids.

**Conclusions:** Allergen exposure induced an elevation in metabolites associated with glycolytic metabolism particularly in the lungs, indicating enhanced activation and increased numbers of immune cells. Notably, inhibition of endocannabinoids mitigated these shifts, underscoring its anti-inflammatory efficacy.

## Introduction

Asthma is a multifaceted and chronic disease, with complex characteristics of heterogeneous airway inflammation, irreversible expiratory flow limitation, and bronchial hyperresponsiveness. This condition is frequently accompanied by symptoms such as wheezing, shortness of breath, cough, and chest tightness, which may vary in intensity and occurrence over time (1). However, this description provides only a general overview of asthma, as it is not a homogeneous disease in terms of its natural history, severity, and response to therapy. Asthma manifests with variable clinical presentations, known as phenotypes, and distinct underlying pathophysiological pathways, referred to as endotypes, which account for this heterogeneity (2). To comprehensively evaluate the various clinical phenotypes and underlying endotypes, a multidimensional approach is required, which encompasses clinical, physiological, immunological, and genetic factors. Such an approach would provide a more accurate and thorough understanding of the underlying mechanisms and the heterogeneity of this condition. For example, the transcriptomics studies conducted revealed consistent associations between the upregulation of type 2 (T2) immune pathways, eosinophilic airway inflammation, and airflow limitation in asthma patients (3–5). These patients are now recognized as having T2-“high” asthma. The T2-“high” asthma phenotype was observed in approximately 50% of the study population, and these patients displayed greater allergic reactions and sensitivity to specific antigens, increased eosinophil counts, increased blood immunoglobulin-E (IgE) levels, and higher bronchial hyperreactivity than those with low expression (6, 7). The discovery of the previous phenotype led to the widespread acknowledgement of a non-T2 or T2-“low” asthma phenotype characterized by increased neutrophil counts, higher resistance to inhaled corticosteroids to less controlled symptoms (8). The complexity of asthma presents a challenge in confirming the diagnosis for more than a quarter of patients receiving primary care (9, 10). Therefore, there is a crucial need for biomarkers that can aid in the diagnosis and endotyping of asthma. This is particularly critical given that currently employed biomarkers are primarily invasive and possess high specificity, but low sensitivity for identifying T2-high asthma. Furthermore, no established biomarkers are available for T2-low asthma. As such, there is an urgent requirement for identifying biomarkers that are both effective and non-invasive in aiding the diagnosis and endotyping of asthma.

Endocannabinoids, a group of lipid-signaling molecules, are synthesized from arachidonic acid and conjugated with either ethanolamine or glycerol. These compounds exhibit binding affinity to cannabinoid receptors, CB1 and CB2, as well as other receptors such as transient receptor potential vanilloid receptor 1 (TRPV1) and G protein-coupled receptor 55 (GPR55) (11). Among them, anandamide (AEA) and 2-arachidonyl glycerol (2-AG) are the most extensively studied endocannabinoids, while palmitoyl ethanolamide (PEA) and oleoyl ethanolamide (OEA) are less investigated. The degradation of endocannabinoids is primarily mediated by the enzymatic actions of Fatty Acid Amide Hydrolase (FAAH) and Monoacylglycerol Lipase (MAGL); however, the biochemical process is not solely reliant on these enzymes. In addition to FAAH and MAGL, the degradation pathway involves the participation of other enzymes such as α/β-hydrolase-6 (Abh6), α/β-hydrolase-12 (Abh12), and cyclooxygenase-2 (COX-2) (11). There is evidence to suggest that targeting endocannabinoid system possess therapeutic properties for combating the inflammatory response and hyperreactivity in asthma-afflicted airways. The suppression of fatty acid amide hydrolase (FAAH) or monoacylglycerol lipase (MAGL) enzymes has demonstrated significant improvements in inflammatory and hyperreactivity markers in both eosinophilic phenotype of ovalbumin-induced allergic airway inflammatory asthma model in guinea pigs (12) and neutrophilic phenotype in lipopolysaccharide (LPS)-induced acute inflammatory asthma model in mice (13). Hence, it is plausible to propose that the regulation of lung endocannabinoid levels through targeted modulation of their metabolism could serve as a promising therapeutic approach for various phenotypes of asthma. Moreover, delving into the metabolic alterations of this system during respiratory ailments holds significant potential in unearthing novel biomarkers that may aid in accurate diagnosis and effective management of these conditions.

The intricate system of biochemical reactions, known as cell metabolism, is crucial for furnishing cells with the requisite energy and substrates for cellular structures. Furthermore, immune cell metabolism not only maintains the energy needs of activated cells but also plays a critical role in the synthesis of essential immunological effector molecules. Therefore, comprehending the metabolic alterations in immune cells during activation is fundamental to gain insights into the pathogenesis of inflammatory diseases and to identify potential immune-metabolic pathways to modulate for improved therapeutic outcomes or enhanced biomarker discovery (14, 15). The analysis of metabolites produced in cells during metabolic processes, referred to as metabolomics, is a valuable approach in studying and investigating low-molecular weight constituents. Consequently, metabolite analysis has increasingly been used in various fields of research due to the significant importance of metabolites in biological systems. It is a powerful method that enhances the understanding of various pathological processes that result from changes in metabolic pathways. Metabolomics is highly sensitive to external stimuli, drug exposures, genetic modifications, and disease pathways, thus making it a useful tool in identifying the pathophysiology of diseases, discovering novel therapeutic targets, and developing personalized therapeutic strategies (16, 17).

Previously, we performed a metabolomic analysis to comprehend the metabolic alterations in the lungs and plasma of mice experiencing acute inflammation and to assess the changes in the levels of major metabolites in the lung linked with endocannabinoid metabolism inhibition. Additionally, we identified potential biomarkers that underwent significant changes during acute inflammation and systemic and local FAAH or MAGL inhibitor treatments (16, 17). In the present study, our objectives are to investigate the metabolic changes that occur during allergic airway inflammation in Guinea Pigs, and to explore how these changes differ between acute and allergic airway inflammation in order to identify novel biomarkers for each phenotype of asthma. Moreover, we aim to gain a better understanding of the role of the endocannabinoid system in allergic asthma and identify novel biomarkers that can be utilized to monitor and regulate treatment courses that target the endocannabinoid system. To achieve these goals, we have broadened our metabolomic analysis to include the BAL cells, where numerous inflammatory cells are typically activated, and blood cells, where inflammatory cells are generally in an inactivated state.

## Methods

In this study, male and female adult Dunkin-Hartley guinea pigs, weighing between 350 and 450 g, were used. The animals were subjected to a 12-hour light-dark cycle and were housed in a controlled environment with regulated ventilation and temperature and given unrestricted access to food. All animal procedures were ethically approved by the Hacettepe University Experimental Animals Local Ethics Committee under protocol number 2020/09-01.

### 2.1. Experimental model for allergic airway inflammation

The experimental animals were randomly assigned to two groups: the unsensitized control group and the group with allergic airway inflammation induced by ovalbumin as previously described (12). Briefly, the ovalbumin group received intraperitoneal (i.p.) injections of a mixture containing 150 μg ovalbumin and 150 mg Al(OH)_3_ on the 1st, 4th, and 7th days of the protocol, with a total volume of 1 ml. Conversely, the unsensitized control group received only 150 mg Al(OH)_3_. To ensure proper delivery, the mixture was injected into two separate areas with a volume of 0.5 ml for each injection. Subsequently, sensitized animals were exposed to 0.3% ovalbumin aerosol for 30 min on the 14th day of the protocol, while unsensitized control animals were subjected to the same inhalation procedure but received only PBS. After 24 h, the guinea pigs were euthanized with high-dose anesthesia (ketamine/xylazine), and experiments and sample collections were performed (Figure 1A).

**Figure 1:**
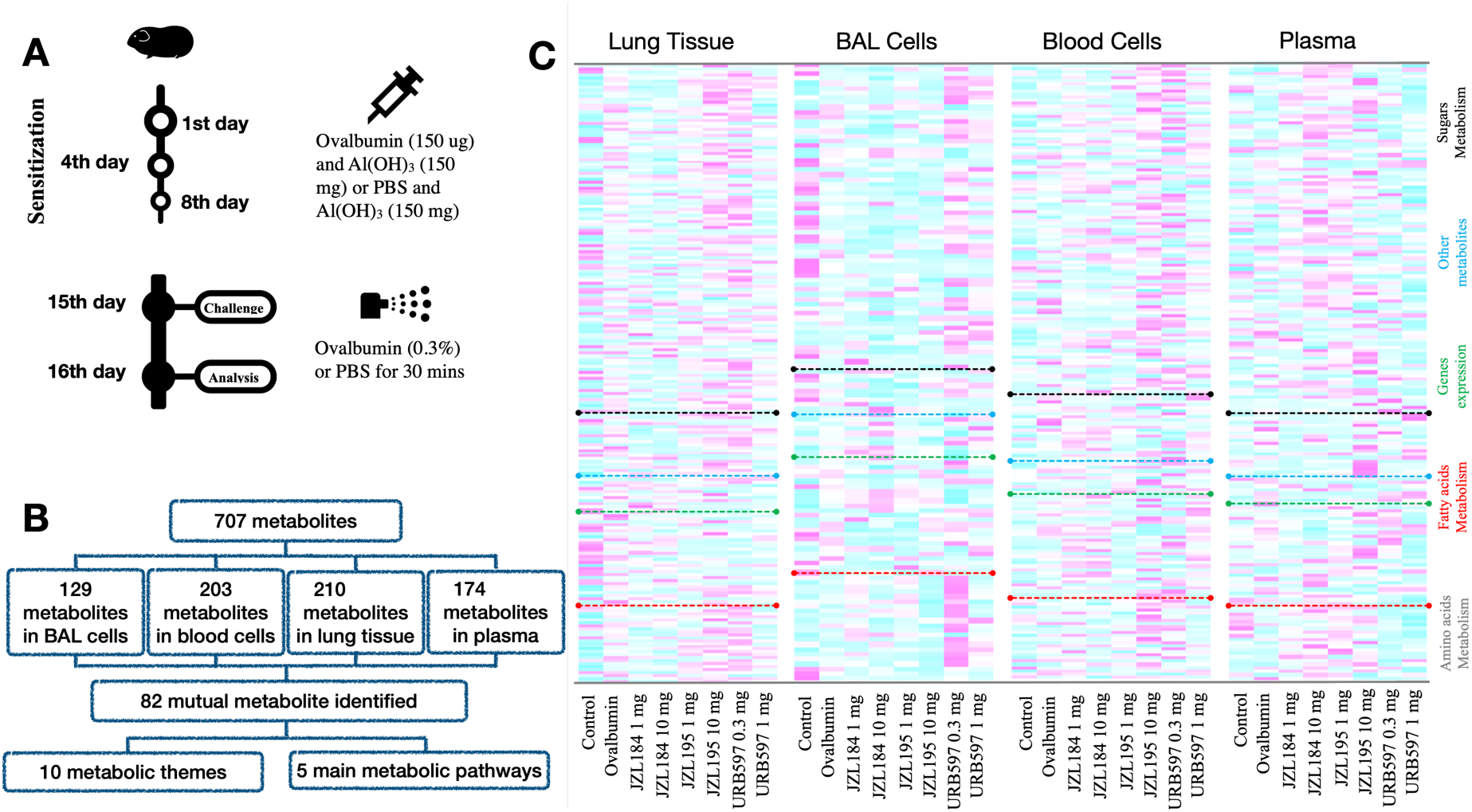
Illustration for the study protocol (A) and the detected metabolite both in numeric flowchart (B) and metabolic heatmap (C).

### 2.2. Treatment with FAAH and MAGL inhibitors

FAAH and MAGL inhibitors applied by i.p. route in a solvent comprising of a mixture of physiological saline, ethanol, DMSO, and Tween80 (17:1:1:1). FAAH inhibitor URB597, MAGL inhibitor JZL184, and FAAH/MAGL dual inhibitor JZL195 are used. The treatments were administered at doses of 0.3 mg/kg and 1 mg/kg for URB597, and 1 mg/kg and 10 mg/kg for JZL184 and JZL195, respectively.

### 2.3. BAL fluid and blood isolation

During the experiment, guinea pigs were anesthetized with ketamine/xylazine to facilitate the cannulation of their trachea. Subsequently, a volume of 3 ml of phosphate-buffered saline (PBS) was instilled into the lungs through the cannula in a slow manner and withdrawn three times. The samples obtained through this procedure were BAL samples. Blood was isolated by cardiac puncture. Blood samples were centrifuged at 1000 g and the plasma was isolated and preserved at −20°C for further analysis. Later, BAL and blood samples were subsequently incubated with lysis buffer for 30 seconds. The samples were then washed with PBS to remove red blood cells and centrifuged to obtain a pellet of cells which were preserved at −80°C for subsequent analyses.

### 2.3. BAL inflammatory cells count

The cellular count procedure involved the incubation of cells with lysis buffer for a duration of two minutes, followed by PBS washing to eradicate red blood cells. The samples were then prepared by applying cytospin slide spraying and subsequently subjected to staining with Giemsa and PAP. An impartial assessment of approximately 300 cells from each sample was carried out by a blinded observer, whereby lymphocytes, eosinophils, neutrophils, and macrophages were counted. The count was conducted by analyzing cells in arbitrary regions.

### 2.4. Sample preparation for metabolomic analysis

The methodology for the preparation of samples and the gas chromatography-mass spectrometry (GC-MS) instrumental conditions were adapted from our previous studies (17). In brief, the lung tissue and BAL and blood cells samples were homogenized by a mortar that was cooled with liquid nitrogen, and 50 μg of fine powder was obtained. Subsequently, the powder was transferred into 1.5 mL microcentrifuge tubes and subjected to extraction with a methanol:water (8:1, v/v) mixture at ambient temperature. For plasma samples, 20 μL plasma were transferred into an Eppendorf tube and extracted using same solution used for tissue. After the extraction, the tissue samples were centrifuged and 200 μL of supernatant was collected in an Eppendorf tube. Then, the samples were completely dried using a vacuum concentrator. Next, the tubes were subjected to methoxyamination and derivatization using N-methyl-N-trimethylsilyltrifluoroacetamide (MSTFA, Thermo Fischer) with 1% trimethylchlorosilane (TMCS, Thermo Fischer). Finally, the samples were analyzed using a Shimadzu GC-MS QP2010 Ultra system.

### 2.5. Metabolomic data analysis

The data processing pipeline involved several steps, including data deconvolution, peak alignment, and normalization, which were carried out using MS-DIAL. The resulting data matrix was screened to exclude metabolite traits with more than 50% missing values. For the remaining missing values in the data matrix, an imputation strategy was applied using the half value of the smallest concentration in the corresponding metabolite group.

The final data matrix was imported into SIMCA-P+ (version 13.0; Umetrics, Umeå, Sweden) for multivariate analysis. Principal component analysis (PCA) was performed to examine the homogeneity of the data, identify any groupings, outliers, and trends, while partial least squares differentiation analysis (PLS-DA) was used to classify the data into different groups, simplify interpretation, and identify potential biomarkers. The variable importance in projection (VIP) graphs were created to distinguish the most important metabolites for the stratification of the groups.

In the examination of metabolic pathways, metabolites were classified based on their functional roles in biological metabolic reactions (refer to Table S1 for the classifications), delineating ten principal metabolic pathways depicted as bar graphs in Figure 3. Moreover, they were stratified according to their participation as either input or output products within five fundamental metabolic pathways, specifically amino acid metabolism, fatty acid metabolism, sugar metabolism, gene expression, and other miscellaneous pathways (refer to Table S1 for the classifications), presented as a heatmap in Figure 1.

To assess the significance of observed distinctions, statistical comparisons of mean expression levels per metabolite among the groups were conducted using the one-way ANOVA test, Mann-Whitney test, and Student’s unpaired t-test. Statistically significant differences were identified when *P* < 0.05. Subsequently, all data were depicted as mean ± standard error. All visualizations were generated utilizing RStudio software (version 2023.12.1 Build 402 © Posit Software PBC).

## Results

A metabolomics study using gas chromatography-mass spectrometry (GC-MS) was conducted on guinea pigs to investigate the impact of fatty acid amide hydrolase (FAAH) and monoacylglycerol lipase (MAGL) inhibition on the metabolomic profile of allergic airway inflammation. The resulting data from GC-MS analysis were subjected to multivariate analysis techniques, including principal component analysis (PCA) and partial least squares-discriminant analysis (PLS-DA), as well as fold-change heatmaps, normalized peak area bar graphs, and fold-change correlation analysis to extract meaningful information. The GC-MS-based metabolomics approach enabled the identification of a total of 707 metabolites across lung (210), BAL cells (129), plasma (174), and blood cells (203) samples, of which 82 were detected in all four sample groups (Figure 1B) and presented individually as bar graphs in (Supp. Figures 5-11).

In metabolomic analyses, pairwise comparisons of metabolomic profiles were conducted initially to assess the impact of the disease model on the metabolomic profiles across diverse matrices. Upon examination of Partial Least Squares Discriminant Analysis (PLS-DA) plots representing the control and ovalbumin groups, a statistically significant distinction was observed across all matrices between the groups (Figure 2). Evaluation of the R2 (the proportion of variance explained by a component) and Q2 (the proportion of total variation predicted by a component) values associated with the applied PLS-DA methodology revealed that the PLS-DA model, indicative of discrimination, exhibited stability and demonstrated robust predictive capability (>0.5).

**Figure 2.**
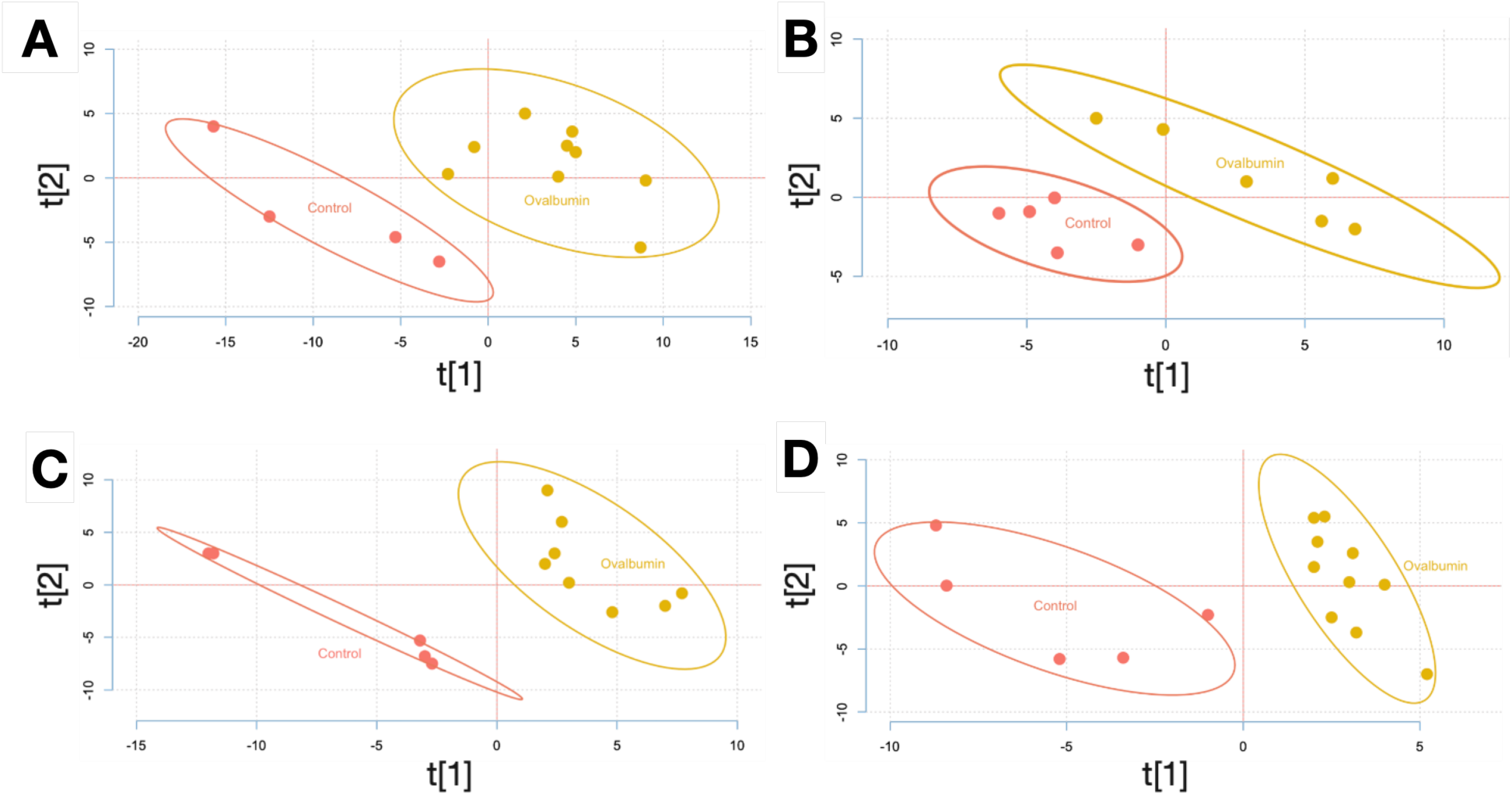
PLS-DA score plots A) lung (R^2^:0.58, Q^2^:0.53), B) BAL cells (R^2^:0.83, Q^2^:0.78) C) blood cells (R^2^:0.48, Q^2^:0.86), and D) plasma (R^2^:0.63, Q^2^:0.74).

The present study also investigated the impact of JZL184, JZL195, and URB597 treatments administered at varying doses on the metabolomic profile of guinea pigs. This analysis was conducted utilizing PLS-DA graphs, focusing on lung tissues (Supp. Fig. 1), bronchoalveolar lavage (BAL) cell samples (Supp. Fig. 2), plasma (Supp. Fig. 3), and blood samples (Supp. Fig. 4). Notably, in lung tissue, plasma, and blood cells, the higher dose of JZL184 (10 mg/kg) exhibited more pronounced distinctions from the ovalbumin group compared to the lower dose (1 mg/kg). However, in BAL cells, no significant variance was observed between the treatment groups and the ovalbumin group. Similarly, JZL195 and URB597 demonstrated analogous patterns to JZL184, with the exception that in lung tissue, the lower dose of JZL195 (1 mg/kg) exhibited greater differentiation from the ovalbumin group than the higher dose (10 mg/kg). Meanwhile, URB597 showed a comparable outcome in blood cells.

In order to attain a more comprehensive evaluation of the results derived from the metabolomics analysis, metabolites exhibiting disparities across groups were categorized based on their role as either input initiators or output by-products within specific metabolic pathways. Subsequently, a heat map was constructed encompassing five principal pathways: amino acids, fatty acids, sugars, gene expression products, and miscellaneous metabolites (Figure 1C). The heat map illustrated that exposure to ovalbumin induced discernible alterations in metabolites within lung and bronchoalveolar lavage (BAL) cells, whereas treatment groups displayed more generalized metabolic changes evident in plasma and blood cells.

Subsequently, to quantify the observed alterations in our analysis, the metabolites were reclassified into distinct categories, including amino acids or those associated with amino acid metabolism, unsaturated or saturated fatty acids, sugar metabolism, glycolysis, the Krebs cycle, energy metabolism, lipid metabolism, nucleotides, kinase residues, and other metabolites. These various metabolic categories were then juxtaposed between control, ovalbumin-exposed, and treatment groups (Figure 3). Notably, exposure to ovalbumin exclusively perturbed the metabolic profile within lung and BAL cells compared to the control group. This perturbation was characterized by an augmentation in inputs crucial for energy metabolism, such as amino acids and fatty acids. Furthermore, disturbances in glycolytic metabolism and metabolites associated with the Krebs cycle were also evident. Moreover, metabolites implicated in activation and signalling pathways, such as lipid metabolism, nucleotides, and kinase residues, exhibited sharper increments subsequent to ovalbumin exposure compared to the control group, particularly within BAL cells and to a lesser extent in lung tissue.

**Figure 3.**
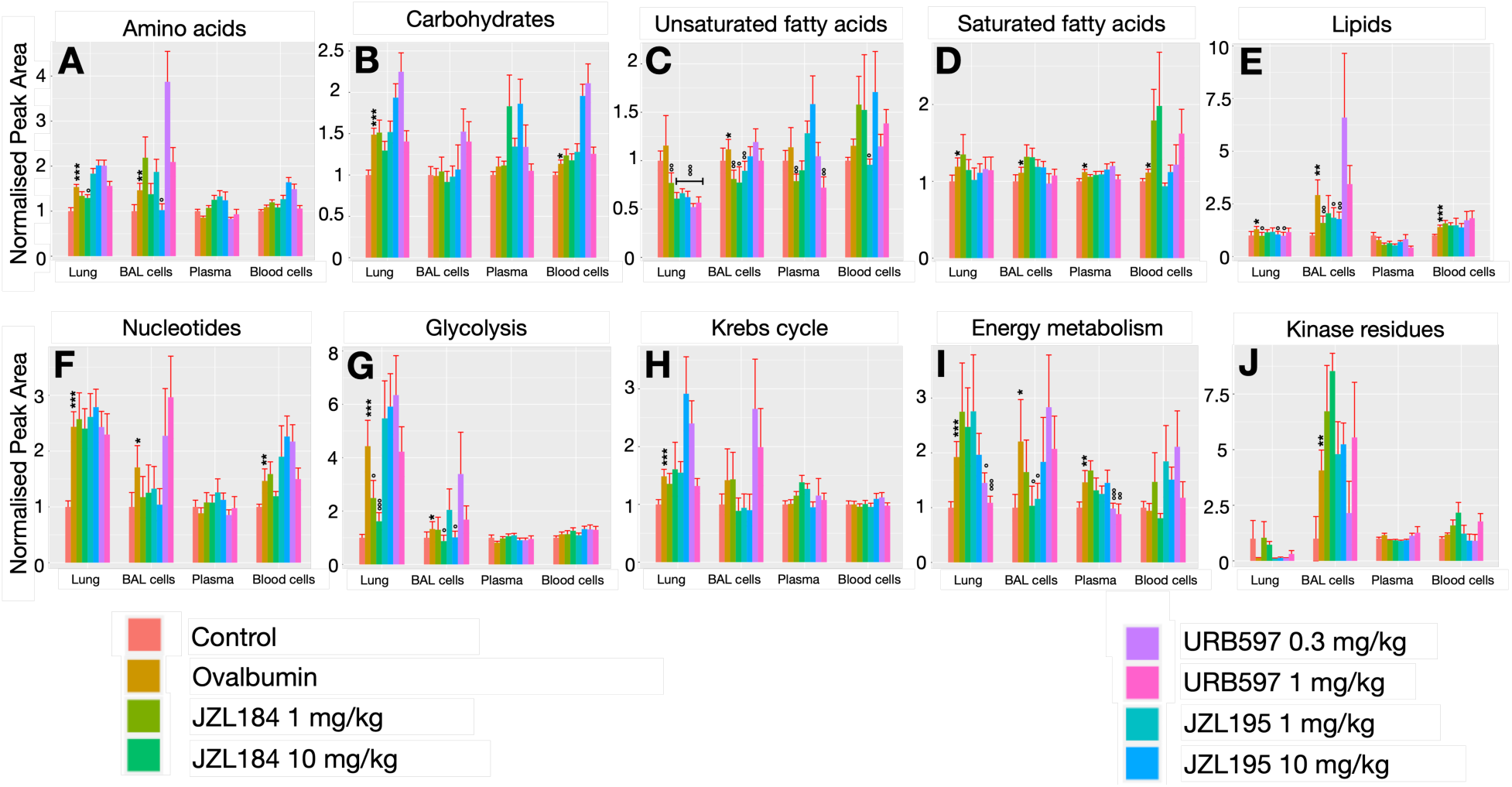
illustrates the alterations in metabolite profiles within the ovalbumin and treatment cohorts relative to the control group. The depicted changes are presented through bar graphs representing various metabolic pathways: (A) amino acid metabolism, (B) carbohydrate metabolism, (C) unsaturated fatty acid metabolism, (D) saturated fatty acid metabolism, (E) bioactive lipid metabolism, (F) nucleotide metabolism, (G) glycolysis metabolites, (H) Krebs cycle metabolites, (I) general energy metabolism, and (J) kinase residues metabolites. Asterisks (*) denote statistically significant differences from the control group, while (**°**) signify distinctions from the ovalbumin group. The significance levels are indicated as follows: * p < 0.05, ** denotes p < 0.01, and *** denotes p < 0.001.

The distinctive alterations observed exclusively in lung and bronchoalveolar lavage (BAL) cells subsequent to allergen exposure, as opposed to blood or plasma, prompted further investigation to elucidate the underlying rationale for these changes. Consequently, a correlation analysis was conducted for individual amino acids (depicted in Figure 4) and fatty acids (illustrated in Figure 5) metabolites subsequent to ovalbumin exposure across lung, BAL cells, blood cells, and plasma. A robust positive correlation among amino acid metabolites was discerned in BAL cells, and to a lesser extent in lung tissue, but not in plasma or blood cells, indicating heightened activity in protein breakdown and synthesis. Conversely, fatty acid-related metabolites exhibited significant positive and negative correlations exclusively in lung tissue, with no such correlations evident in the other specimens, suggesting an upsurge in the on-demand synthesis of inflammatory lipid molecules.

**Figure 4.**
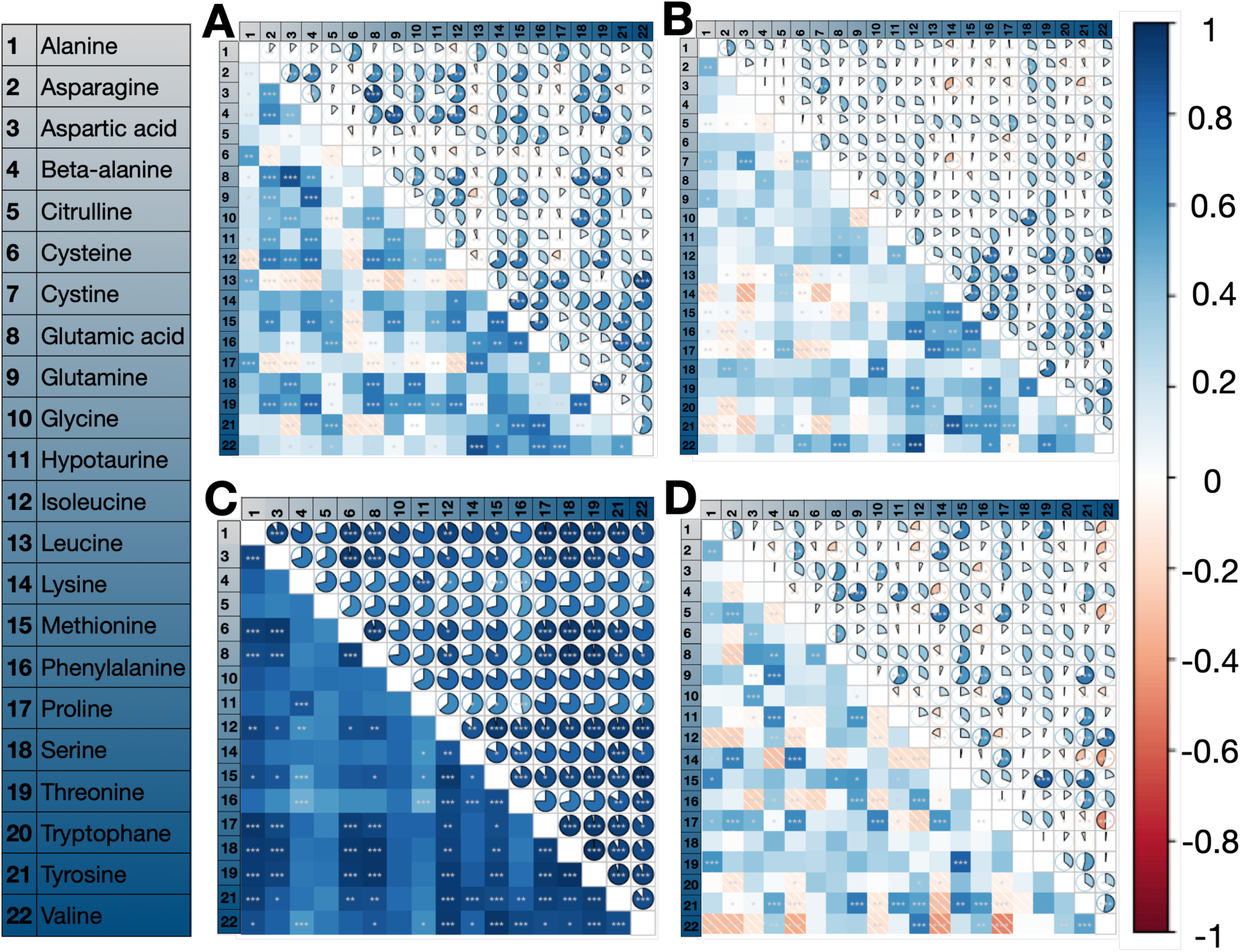
displays a heatmap illustrating the Pearson correlation coefficient outcomes derived from the correlation analysis conducted among amino acids present in the (A) lung tisse, (B) plasma, (C) bronchoalveolar lavage (BAL) cells, and (D) blood cells. The significance levels are denoted as follows: * for P < 0.05, ** for P < 0.01, and *** for P < 0.001.

**Figure 5:**
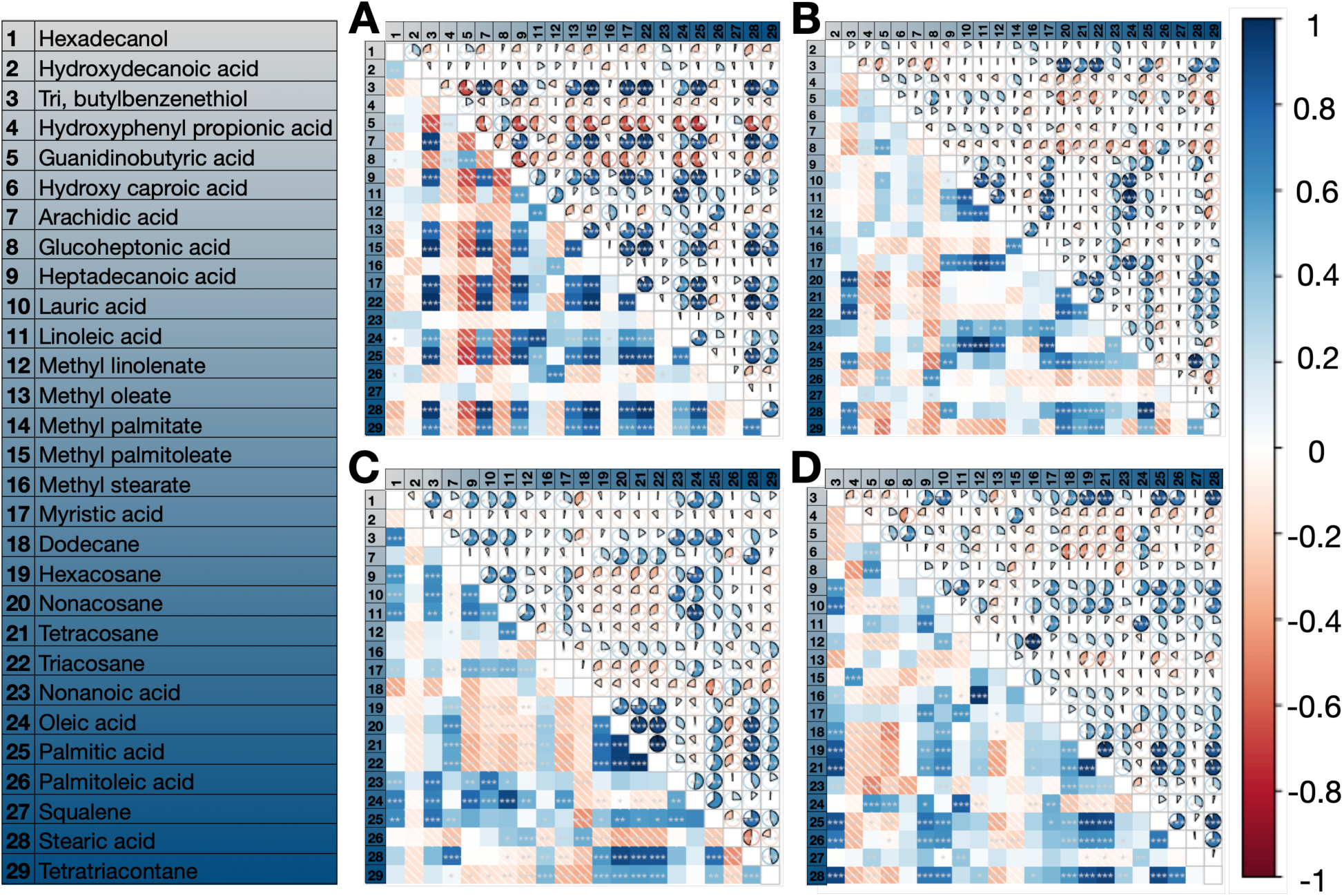
displays a heatmap illustrating the Pearson correlation coefficient outcomes derived from the correlation analysis conducted among fatty acids present in the (A) lung, (B) plasma, (C) bronchoalveolar lavage (BAL) cells, and (D) blood cells. The significance levels are denoted as follows: * for P < 0.05, ** for P < 0.01, and *** for P < 0.001.

**Figure 6:**
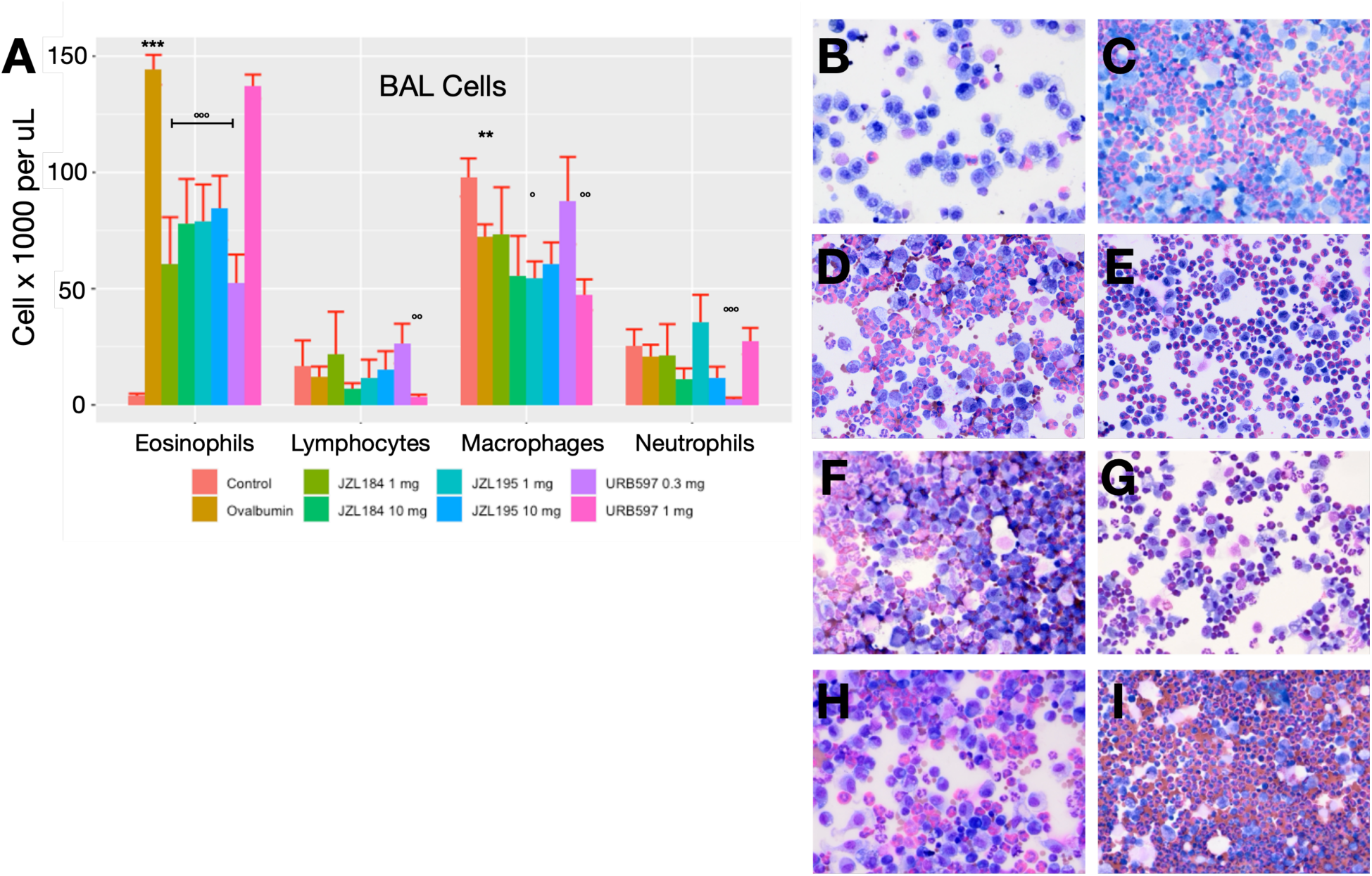
displays the quantification of BAL (Bronchoalveolar Lavage) cell counts, depicted both in (A) bar graph format and (B-I) visual representations that encompasses various treatment conditions, including (B) control, (C) ovalbumin, (D) JZL184 1 mg/kg (E) JZL184 10 mg/kg, (F) JZL195 1 mg/kg (G) JZL195 10 mg/kg, (H) URB597 0.3 mg/kg, and (I) URB597 1 mg/kg. The symbol ‘*’ signifies a statistically significant difference compared to the control group, while (°) denotes a difference from the ovalbumin group. The significance levels are denoted as follows: * for P < 0.05, ** for P < 0.01, and *** for P < 0.001.

In contrast, the suppression of endocannabinoid metabolism resulted in a significant systemic alteration in energy metabolites, including carbohydrates, amino acids, and fatty acids, when compared to the control group. Furthermore, inhibiting MAGL, more than inhibiting FAAH, exhibited greater efficacy in mitigating the observed pathway modifications following ovalbumin exposure in lung and BAL cells. These alterations encompassed amino acid metabolism, carbohydrate metabolism, glycolysis, fatty acid metabolism, the Krebs cycle, energy metabolism, as well as activation and signalling pathways related to lipid metabolism, nucleotide metabolism, and kinase activation (Figure 3).

### The airway and lung histopathology

In order to validate the alterations identified in the metabolomic analysis during allergic inflammatory processes, our objective was to conduct a pathological examination to enumerate bronchoalveolar lavage (BAL) cells. The present investigation documented a notable increase in eosinophil count in BAL subsequent to ovalbumin exposure when compared to the unsensitized PBS-exposed cohort. This escalation was effectively mitigated in all treatment cohorts except for the URB597 (1 mg/kg). No discernible variance was observed in lymphocyte and neutrophil counts subsequent to ovalbumin exposure.

## Discussion

The pathophysiology of asthma involves various signaling molecules and cells. Eosinophilic asthma is an asthma phenotype that exhibits a high prevalence of eosinophils in both airways and peripheral circulation. Animal models have been developed to investigate this phenotype, which typically manifests in conjunction with allergic reactions. Guinea pigs sensitized to ovalbumin have been widely used in such models due to their high sensitivity to allergens. However, the metabolic analysis of these models is only recently gaining popularity. Martin et al. (18) conducted a metabolic analysis of sensitized guinea pigs’ plasma and identified several potential metabolites that could be associated with eosinophilic asthma. Nevertheless, studying the plasma alone is insufficient since it provides a generalized sample and disregards the crucial inflammatory processes and reactions that occur in the local inflammation area. Therefore, we conducted a more comprehensive metabolomic analysis that examined not only plasma but also lung tissue, blood cells, and BAL cells from sensitized animals. In fact, our plasma analysis yielded comparable outcomes to Martin et al.’s findings concerning changes in tryptophan, taurine, and other potentially NO and ROS-related metabolites. Furthermore, using an extensive analysis, we have elucidated alterations in the energy pathways and discerned potential biomarkers that are specifically associated with the eosinophilic asthma phenotype.

Metabolism represents a highly intricate and multifaceted network of biochemical reactions, which are orchestrated with the purpose of endowing the cell with the appropriate molecules at the appropriate moment. To expound, the intricate process of metabolism involves the breakdown of glucose, fatty acids, and amino acids with the intention of generating both energy in the form of ATP and metabolites that serve as the essential building blocks of various cellular pathways and components (14). To meet the energy demands of cellular processes, glucose undergoes a complex enzymatic process called glycolysis, consisting of multiple sequential reactions that yield two molecules of pyruvate and two molecules of ATP. Following glycolysis, pyruvate enters the mitochondrion and undergoes further oxidative metabolism in the Krebs cycle where much more ATP molecules are generated. This cycle can also be supplemented through the introduction of acetyl-CoA from fatty acid oxidation or glutamate from amino acid metabolism at specific reaction points (19). Cells that exhibit a high rate of proliferation and immune cells that are activated can convert the pyruvate produced during glycolysis into lactate. This metabolic pathway, known as lactate fermentation, is characterized by a lower energetic efficiency compared to oxidative phosphorylation. Lactate fermentation involves only a partial oxidation of the carbon atoms present in glucose, thereby limiting the amount of energy generated per glucose molecule. However, this process enables a faster production of ATP, and is not constrained by oxygen availability. Hence, under conditions of high energy demand, such as during rapid cell division or in response to an immune challenge, cells can resort to glycolysis as a means of quickly generating ATP to meet their metabolic needs (20). This selective catabolism of glucose to lactate is known to disrupt the Krebs cycle, as a consequence of inadequate pyruvate supply to the mitochondria. This impairment leads to specific arrest points in the Krebs cycle, which consequently results in the accumulation of intermediary metabolites (14). For example, research on macrophages stimulated with LPS has revealed that these arrest points occur downstream of citrate and succinate (21, 22). The present study reveals a clear manifestation of a phenomenon, wherein exposure to ovalbumin triggers an increase in metabolite associated with the glycolysis metabolism in the lungs, but not in plasma. This implies a heightened activation and numbers of immune cells and other energy-demanding cells. In addition, we observed a significant increase in citric acid levels in the lung tissue of the ovalbumin-treated group compared to the control group, suggesting a disrupted Krebs cycle in the lung. This observation is further supported by the finding that metabolites related to Krebs cycle dysfunction were elevated in the lung tissue and BAL cells, but not in the plasma or blood cells. Moreover, our findings indicate that MAGL inhibition, but not FAAH inhibition, tends to hinder this metabolic shift in the lungs in a dose-dependent manner. This effect is plausibly linked to MAGL inhibition’s impact on cell migration into the lung, given that our current investigation found a significant reduction in the number of inflammatory cells in BAL, as well as a decrease in the total cell count in BAL from our prior study (12).

In mice treated with HDM, RNA-sequencing analysis revealed the highest upregulation of the gene encoding for the fatty acid desaturating enzyme stearoyl-coenzyme A desaturase compared to those treated with PBS. Furthermore, a gene cluster related to lipid biosynthesis processes also demonstrated upregulation (23). Our study found that levels of saturated fatty acids slightly increased in the ovalbumin group compared to the control group in lung, plasma, BAL, and blood cells. Additionally, lipid metabolism slightly increased in the lung and dramatically increased in activated BAL cells. Thus, our findings support Nishimura et al.’s results that suggest lipid metabolism is a promising therapeutic target for allergic airway inflammation.

In this context, it was observed that the lipid glyceryl oleate exerted an impact on the immune function of dendritic cells, leading to a downregulation of the expression of genes associated with Th2 response (24). Our investigation revealed a significant reduction in the levels of methyl oleate in the ovalbumin-treated group as compared to the control group. Notably, the inhibition of MAGL not only counteracted this reduction but also attenuated the accumulation of inflammatory cells in BAL.

Imbalances in amino acid metabolism have been shown to impair lung function in asthmatics, with particular attention paid to glutamine and tryptophan metabolites, including urea, ornithine, and citrulline (25). These metabolites have been found to play a crucial role in altering the balance between regulatory T cells (Tregs) and T helper 17 (Th17) cells, as well as in the production of OVA-specific immunoglobulin E (IgE), interleukin-5 (IL-5), and lung inflammation (26). Ornithine metabolism has been suggested as a possible treatment for allergic asthma (27), while citrulline supplementation has been shown to improve lung function in obese asthmatics (25). In our study, the levels of both ornithine and citrulline significantly decreased after ovalbumin exposure in BAL cells, highlighting the potential of these metabolites as targets for treatment or biomarkers for allergic asthma. This finding is of particular importance, given that the levels of amino acids in BAL cells are strongly correlated, unlike other tissues and cells in our study. However, it is important to note that this correlation may be due to the increased expression of proteins and cytokines in these cells, and further research is necessary to better understand this outcome.

In lung myofibroblasts, the activation of transcription factor 4 leads to an increase in the expression of enzymes involved in the serine-glycine synthesis pathway, which results in the excessive deposition of collagen proteins and scarring in idiopathic pulmonary fibrosis. Additionally, the increased glycolytic flux promotes the conversion of the glycolytic intermediate 3-phosphoglycerate into glycine, which is the most abundant amino acid found in collagen protein (28) . In our study, we observed a significant increase in the level of serine-glycine in lung and BAL cells following a three-week protocol of ovalbumin sensitization and exposure. Moreover, the level of 3-phosphoglycerate was dramatically decreased in BAL cells, indicating a potential association between these markers and collagen deposition and remodeling related to allergic airway inflammation. However, it is important to note that our study did not directly measure the level of collagen deposition, and further data are needed to establish this relationship.

In their study, Zhu et al. (29) employed an objective metabolomics method using bronchoalveolar lavage fluid (BALF) samples from 12 patients who underwent bronchial allergen challenge to induce acute inflammatory responses. Their findings indicated a notable increase in saturated fatty acid synthesis and mitochondrial beta-oxidation of saturated fatty acids following allergen challenge. We observed a similar elevation in saturated fatty acid levels in all tissues, including BAL cells, in response to ovalbumin exposure, which is consistent with the results of Zhu et al. (29) Furthermore, bronchial smooth muscle (BSM) in individuals with asthma has been shown to exhibit a shift towards elevated rates of mitochondrial respiration and increased utilization of fatty acids (30). In a previous investigation, the authors reported a marked reduction in stearic and linoleic acid levels in BSM cells of asthmatic patients as compared to non-asthmatic controls. Our research revealed a similar pattern, with reduced levels of stearic and linoleic acid only observed in the lungs of guinea pigs following exposure to ovalbumin. These results support the findings of Esteves et al. and suggest that these fatty acids may serve as potential biomarkers for airway remodeling in individuals with asthma, given their association with the increased consumption of BSM.

Through the application of ultra-high performance liquid chromatography-mass spectrometry, Liu and colleagues (31) conducted a characterization of metabolic profiles in induced sputum samples derived from both healthy participants and individuals diagnosed with asthma. Nicotinamide concentrations in sputum samples were found to predict rate ratios of severe asthma exacerbations. In our own investigation, we observed a significant increase in the concentration of 1-methyl nicotinamide in lung tissue following exposure to ovalbumin. Moreover, the metabolism pathway of nicotinamide demonstrated significant differences between the eosinophilic and neutrophilic asthma phenotypes in the analysis conducted by Liu et al. Furthermore, the metabolism of tyrosine, phenylalanine, taurine, and hypotaurine served as an effective discriminator between the two aforementioned phenotypes. The concentration of tyrosine in the bloodstream has been found to be lower in individuals with asthma who have lower sensitization compared to those with atopic asthma who exhibit higher sensitization (32). In our study, we observed a significant increase in the levels of tyrosine and hypotaurine in eosinophil-rich bronchoalveolar lavage cells, but not in other tissues. Moreover, a noteworthy reduction in the tyrosine concentration was observed in the blood cells of guinea pigs exposed to ovalbumin. Notably, no such alterations were observed in the lung profile of our previous study utilizing neutrophilic acute inflammation in mice (16). Therefore, the altered metabolism of tyrosine and hypotaurine may serve as promising biomarkers for allergic asthma.

In a previous study by Chiu et al. (33), a metabolomic analysis of urine was conducted to compare asthmatic children to age-matched controls. The results indicated that the degradation-metabolism of valine and isoleucine in urine was significantly associated with allergic sensitization for childhood asthma. In the present study, we have observed a decrease in the plasma level of valine and isoleucine subsequent to ovalbumin exposure, suggesting that these two amino acids may serve as promising biomarkers for allergic asthma. In a separate study, Chiu et al. (32) reported a significant correlation between isobutyric acid levels in the blood and high sensitivity in asthma patients, in comparison to those with lower sensitivity. Additionally, glutamine levels were strongly associated with IgE levels. Similarly, our research findings indicate a significant increase in plasma and lung levels of 3-aminobutyric acid in ovalbumin-sensitized animals, along with a marked elevation in glutamic acid levels in the lung tissue. These findings further support the potential of these metabolites as biomarkers for allergic diseases.

Saghatelian et al. (34) established the interaction between the endocannabinoid system and N-acyl taurine lipids, which are primarily regulated by FAAH. Consequently, the inhibition of FAAH would significantly affect the levels of taurine lipid species. Our previous findings have shown that FAAH inhibition resulted in an increase in hypotaurine concentrations in the lungs of mice (17). Our current results align with these previous findings, as both selective and non-selective inhibition of FAAH resulted in elevated levels of hypotaurine in the lung, bronchoalveolar lavage fluid, and blood cells of treated Guinea Pigs. In concordance with prior research, Côté et al. (35) have documented a correlation between elevated 2-arachidonoylglycerol (2-AG) levels and a reduction in high-density lipoprotein-cholesterol levels. Building upon our earlier research where cholesterol levels decreased in mice lung (17), our current findings reveals a marked decline in cholesterol levels across various biological compartments, including guinea pigs lung, plasma, and blood cells, consequent to the inhibition of monoacylglycerol lipase (MAGL).

Previous studies have demonstrated that cannabinoids possess a regulatory effect on pyruvate metabolism. Specifically, in striated muscle cells of CB1-knockout mice, the expression of pyruvate metabolism genes was found to increase (36), while CB1 antagonism was observed to upregulate pyruvate metabolism enzymes (37). In our previous study conducted in mice, we also observed that the inhibition of endocannabinoid metabolism significantly affected a substantial portion of energy uptake and the production of metabolites (17). These findings suggest a strong association between endocannabinoids and energy metabolism. In our current study conducted on guinea pigs, we provide further evidence to support this association, as we observed that FAAH and MAGL inhibition appeared to impact sugar metabolism in inflamed lungs, plasma, blood, and BAL cells.

## Conclusions

Exposure to ovalbumin exclusively modified the metabolic profile within the lungs of guinea pigs, leading to changes in glycolytic metabolism and alterations in nucleotides in both lung tissue and bronchoalveolar lavage cells. Conversely, inhibiting endocannabinoids metabolism prompted a systemic reconfiguration of energy metabolites, including carbohydrates, amino acids, and fatty acids, with more pronounced effects observed in plasma and blood cells. The increase in metabolites induced by allergen exposure was linked to glycolytic metabolism alterations, particularly in the lungs and BAL cells, indicating heightened activation and proliferation of immune cells. Notably, endocannabinoids metabolism inhibition ameliorated these metabolic shifts, highlighting its anti-inflammatory efficacy.

## Supplementary data

**Supplement Figure 1.**
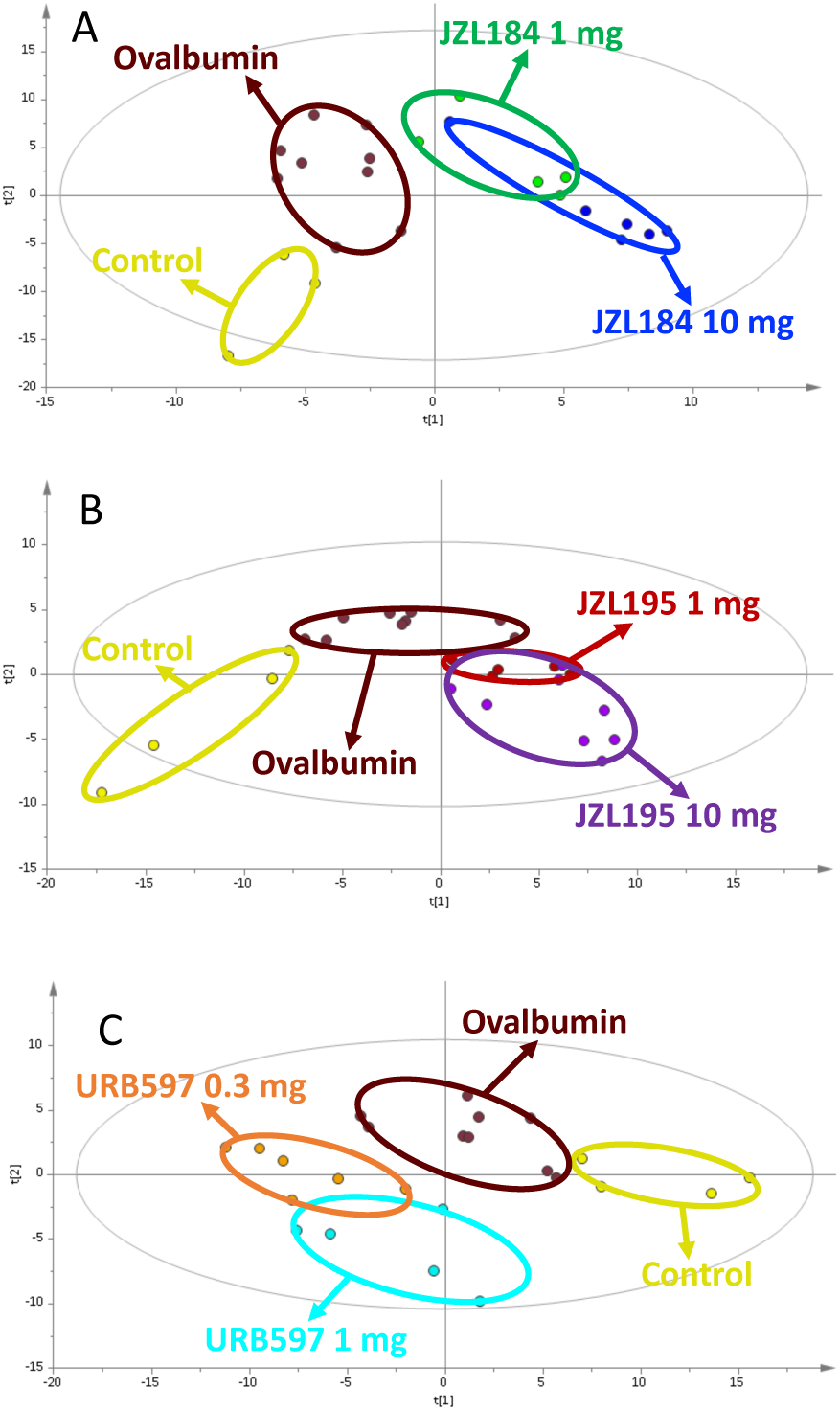
PLS-DA score plots of Lung samples A) JZL184 treatments (R^2^: 0.53, Q^2^: 0.24) B) JZL195 treatments (R^2^: 0.51, Q^2^: 0.26) C) URB597 treatments (R^2^: 0.69, Q^2^: 0.50). For both doses, it was observed that the JZL184 treatment groups showed a different profile from the ovalbumin group, but the 10 mg application differed more from the ovalbumin group than the 1 mg application (Supp. Fig. 1A). It was observed that JZL195 treatment groups showed a metabolomic profile close to the ovalbumin group, but the 10 mg application differed slightly from the ovalbumin group compared to the 1 mg application (Supp. Fig. 1B). It was determined that the URB597 treatment groups showed a different profile from the ovalbumin group for both doses (Supp. Fig. 1C). It was also observed that all treatment groups had a metabolomic profile quite different from the control group.

**Supplement Figure 2.**
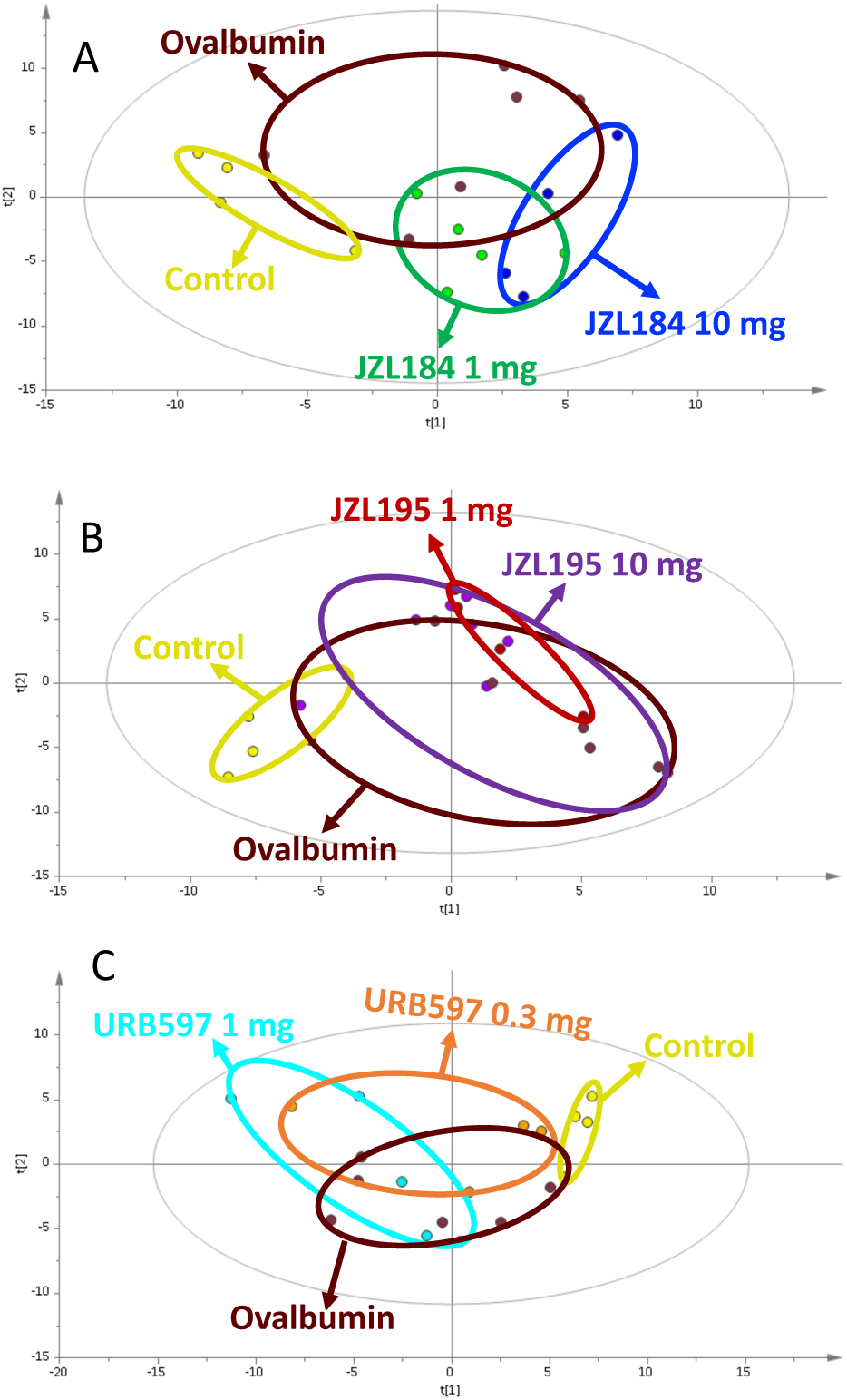
PLS-DA score plots of BAL cell samples A) JZL184 treatments (R^2^: 0.47, Q^2^: −0.02) B) JZL195 treatments (R^2^: 0.57, Q^2^: −0.02) C) URB597 treatments (R^2^: 0.59, Q^2^: −0.16). The effects of JZL184, JZL195 and URB597 treatments on the metabolomic profile in BAL cell samples were examined with PLS-DA graphs (Supp. Fig. 2). However, BAL cell samples did not differentiate the treatment groups from the ovalbumin group to a significant extent, for all treatment applications.

**Supplement Figure 3.**
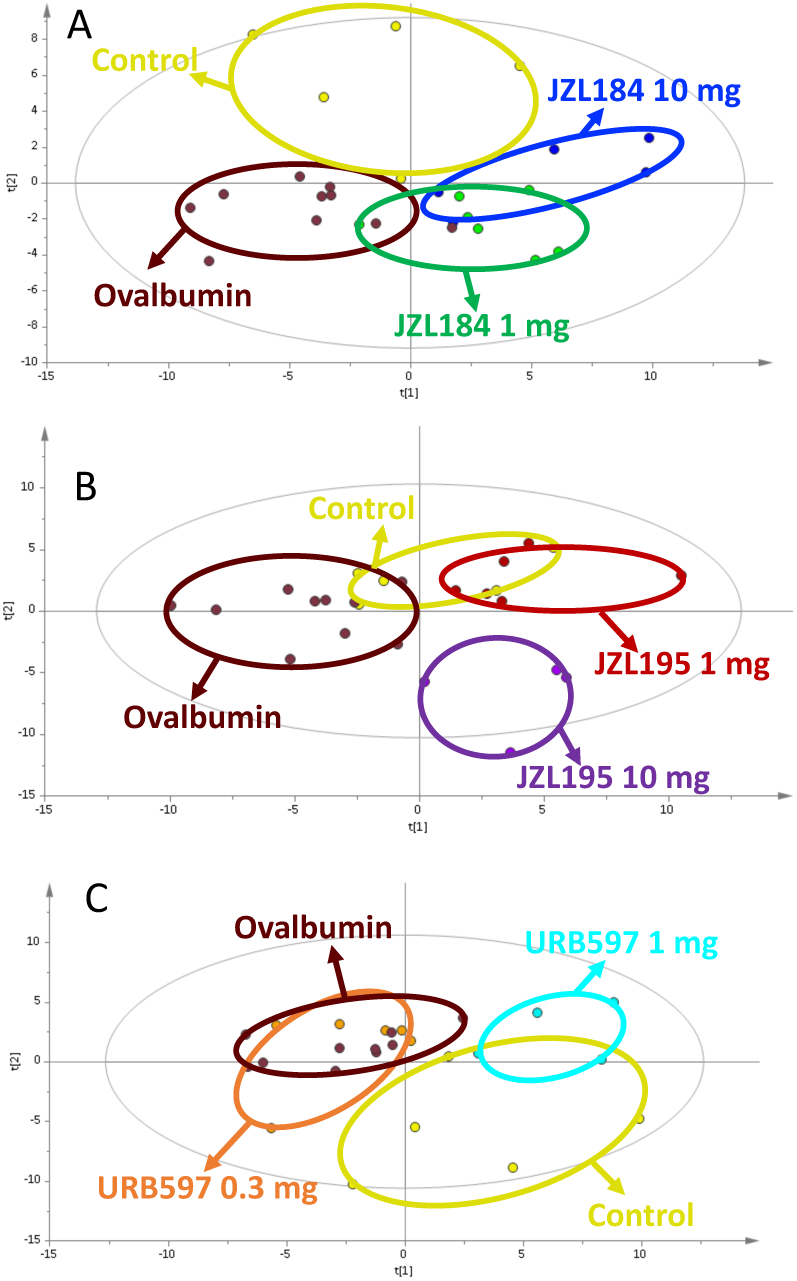
PLS-DA score plots of plasma samples A) JZL184 treatments (R^2^: 0.25, Q^2^: −0.15) B) JZL195 treatments (R^2^: 0.33, Q^2^: 0.21) C) URB597 treatments (R^2^: 0.26, Q^2^: −0.12). The effects of different doses of JZL184, JZL195 and URB597 treatments on the metabolomic profile in the plasma of guinea pigs were examined with PLS-DA graphs (Supp. Fig. 3). For both doses, it was observed that the JZL184 and URB597 treatment groups showed a different profile from the ovalbumin group, but the high dose application differed more from the ovalbumin group than the low dose application (Supp. Fig. 3A and 3C). JZL195 treatment groups appeared to show a significantly different metabolomic profile than the ovalbumin group (Supp. Fig. 3B).

**Supplement Figure 4.**
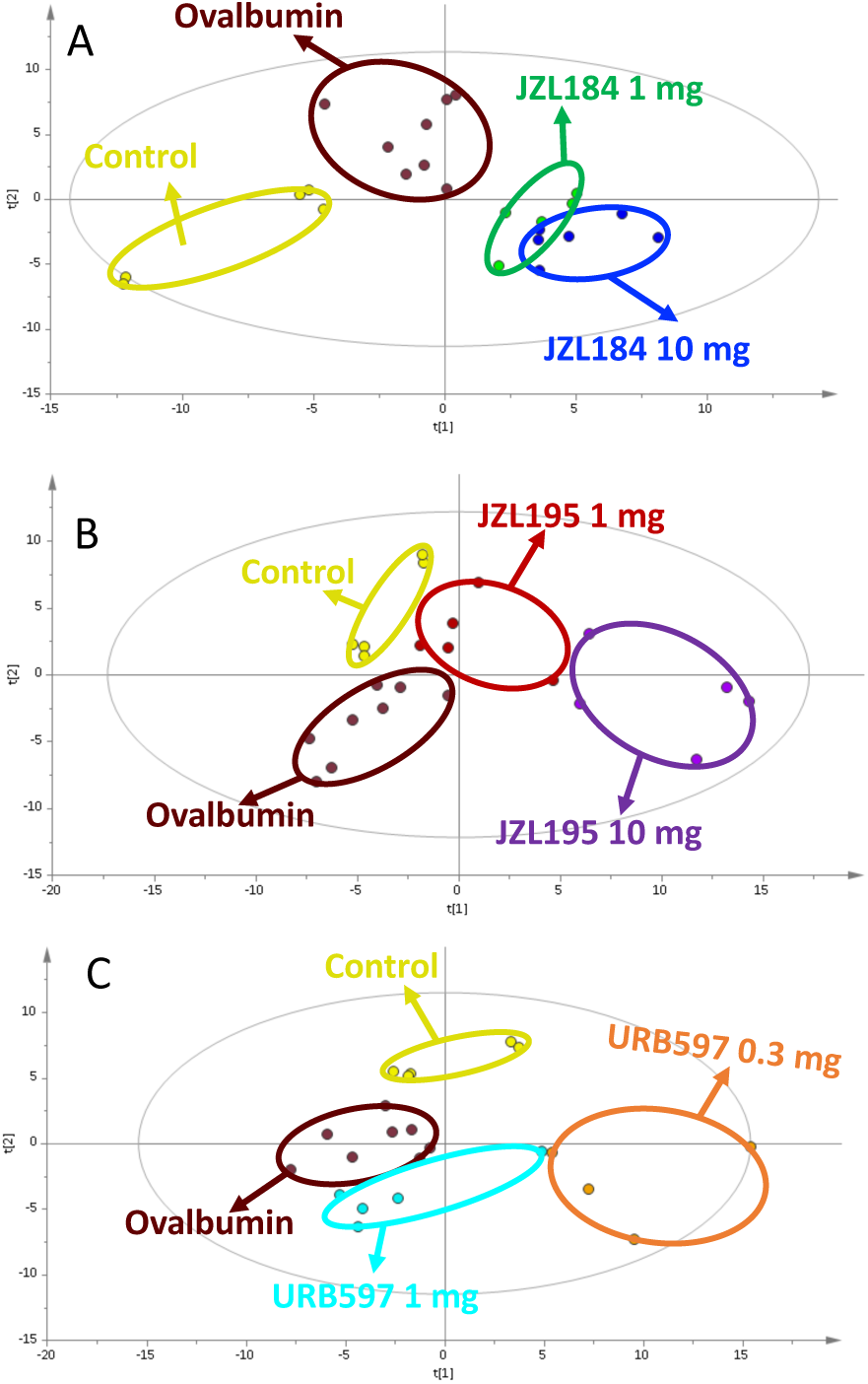
PLS-DA score plots of Blood cell samples A) JZL184 treatments (R^2^: 0.44, Q^2^: −0.32) B) JZL195 treatments (R^2^: 0.51, Q^2^: 0.52) C) URB195 treatments (R^2^: 0.55, Q^2^: 0.12). The effects of two-dose treatments JZL184, JZL195 and URB597 on the metabolomic profile in blood samples of guinea pigs were examined with PLS-DA graphs (Supp. Fig. 4). It was observed that the JZL184 treatment groups showed a different profile from the ovalbumin group, but the 10 mg application differed more from the ovalbumin group than the 1 mg application (Supp. Fig. 4A). In JZL195 treatment groups, it was observed that 1mg application showed a metabolomic profile close to the ovalbumin group, but 10mg application differed more from the ovalbumin group compared to 1mg application (Supp. Fig. 4B). In the URB597 application, while the low-dose treatment showed a different profile from the ovalbumin group, it was determined that the groups had a similar profile in the high-dose application (Supp. Fig. 4C).

**Supplement Figure 6:**
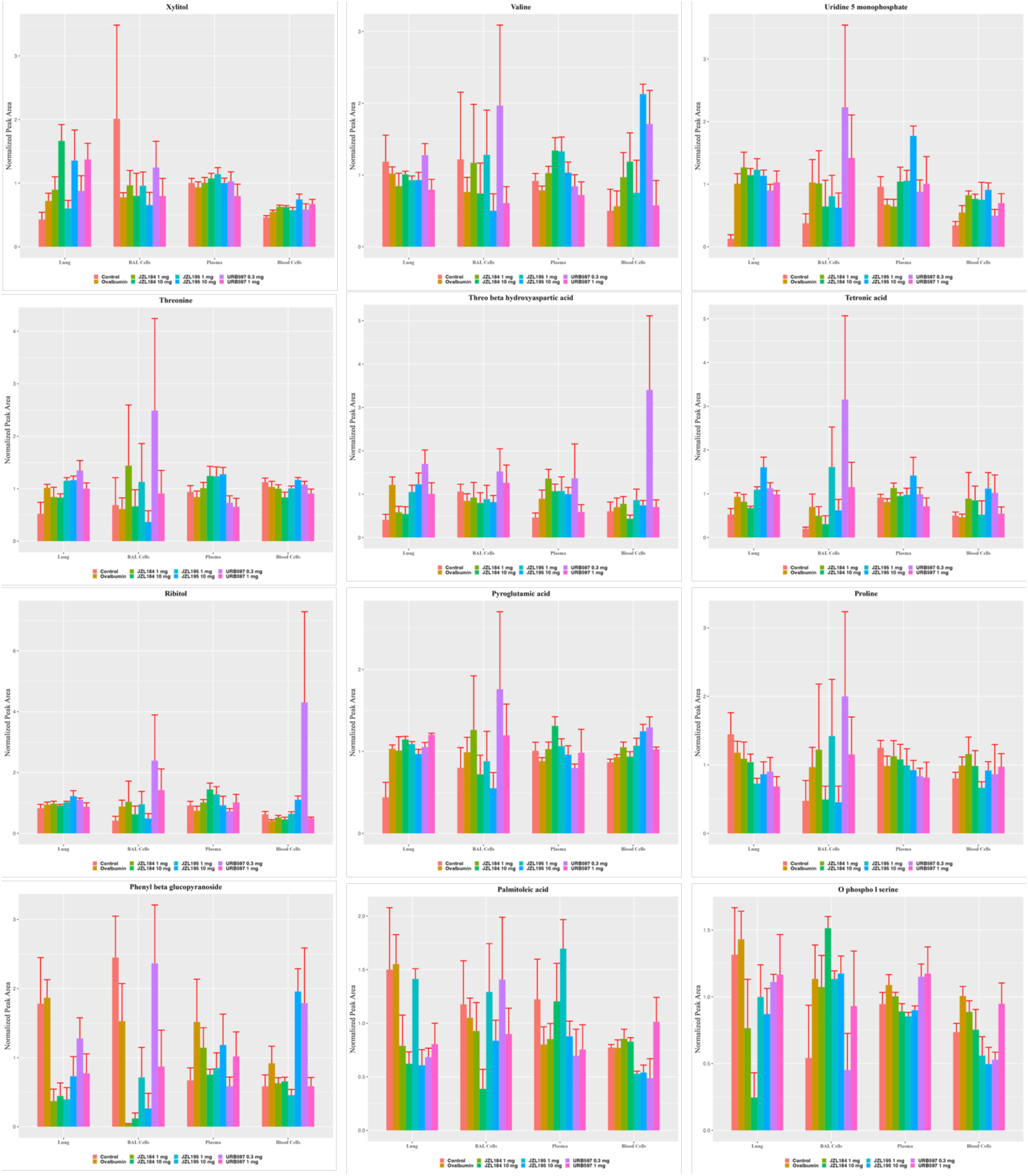
illustrates the alterations in different metabolites within the ovalbumin and treatment cohorts relative to the control group. The depicted changes are presented through bar graphs.

**Supplement Figure 7:**
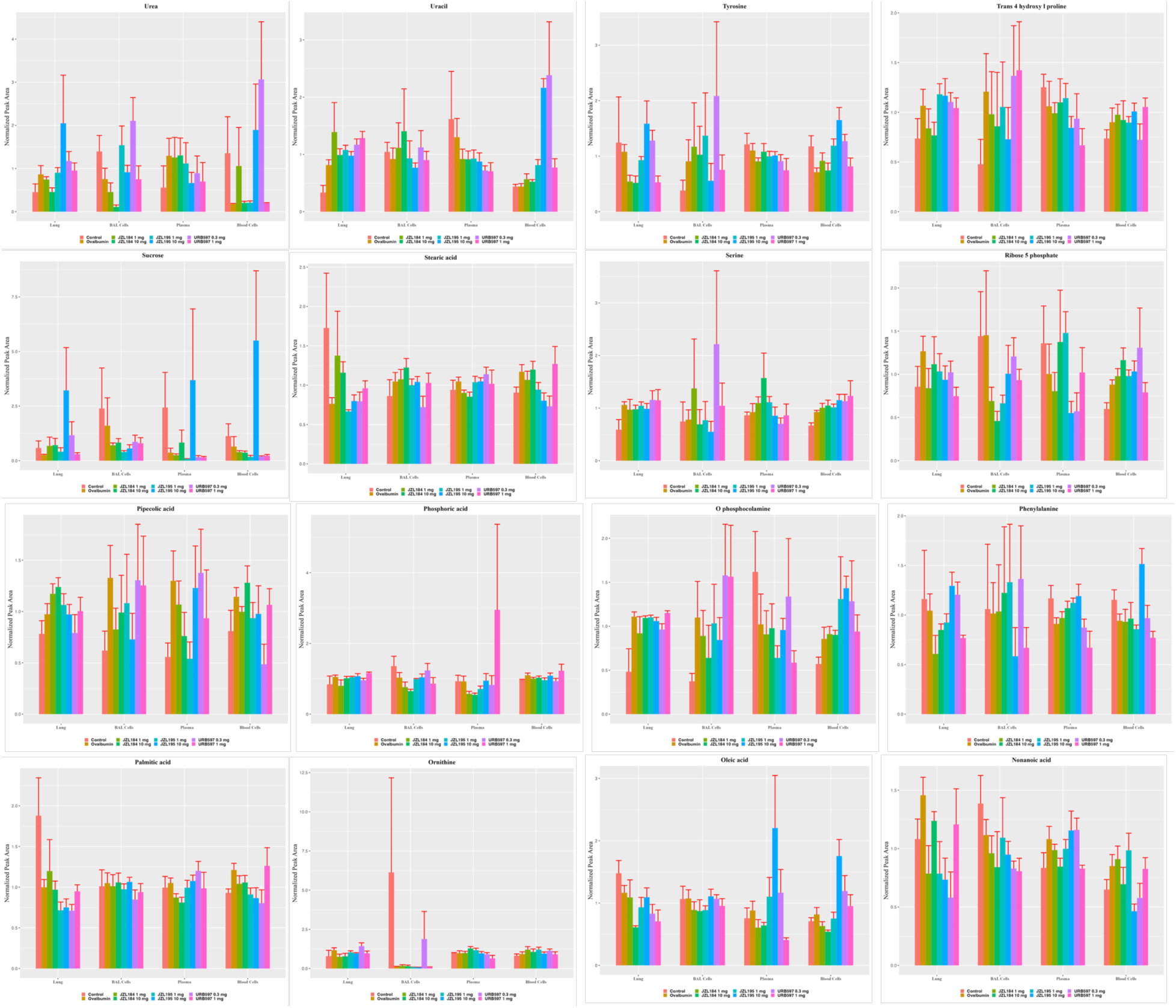
illustrates the alterations in different metabolites within the ovalbumin and treatment cohorts relative to the control group. The depicted changes are presented through bar graphs.

**Supplement Figure 8:**
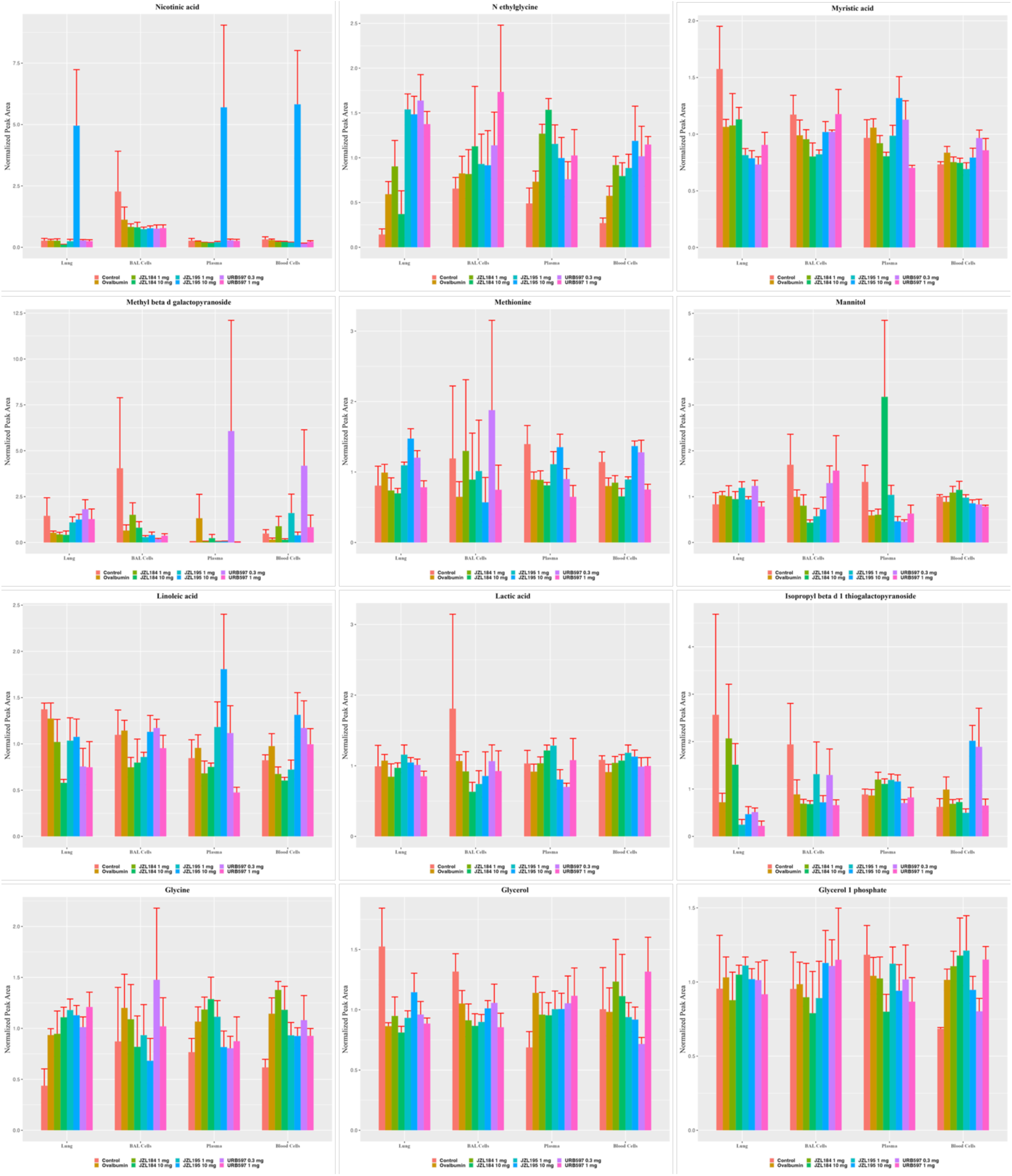
illustrates the alterations in different metabolites within the ovalbumin and treatment cohorts relative to the control group. The depicted changes are presented through bar graphs.

**Supplement Figure 9:**
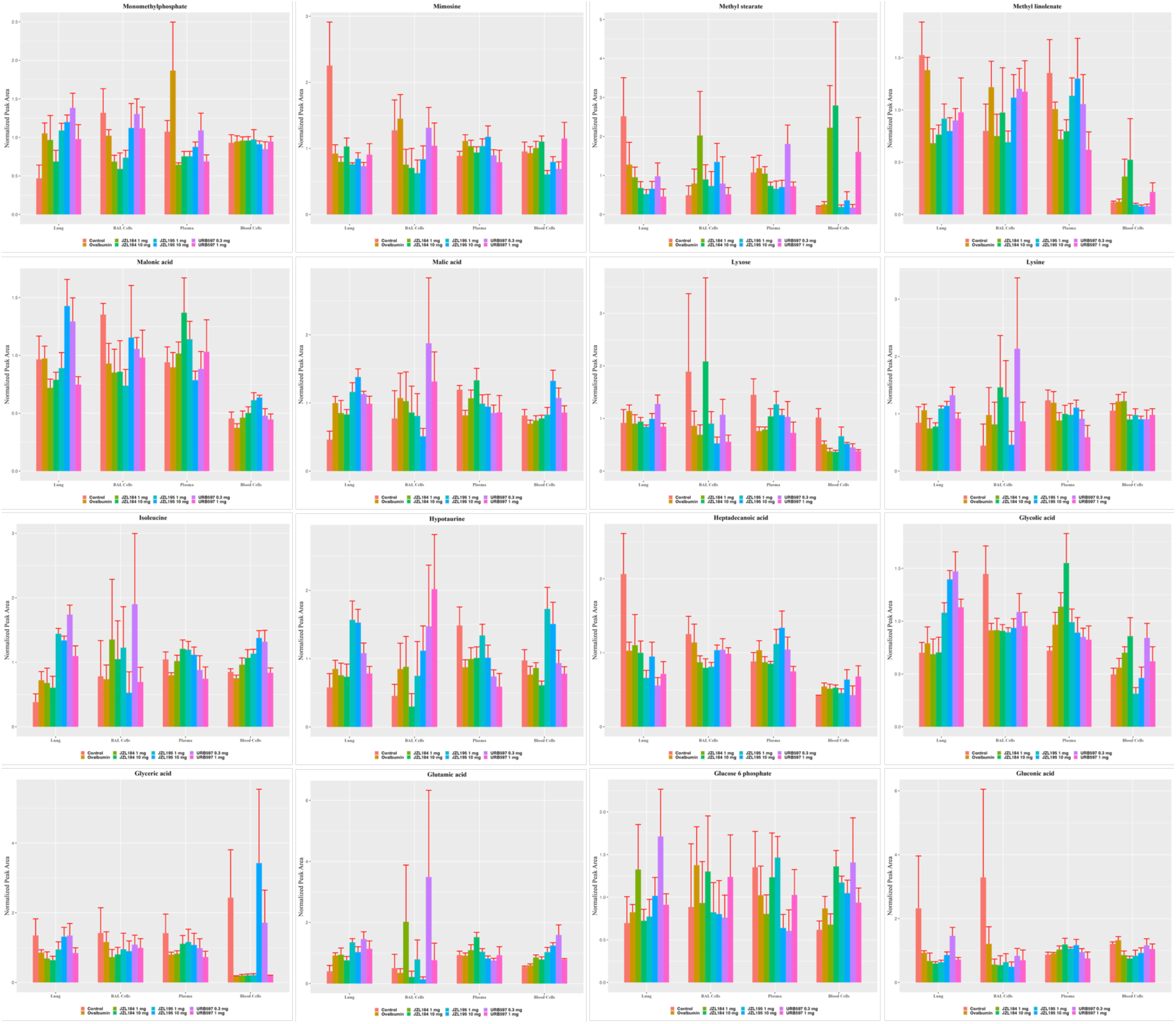
illustrates the alterations in different metabolites within the ovalbumin and treatment cohorts relative to the control group. The depicted changes are presented through bar graphs.

**Supplement Figure 10:**
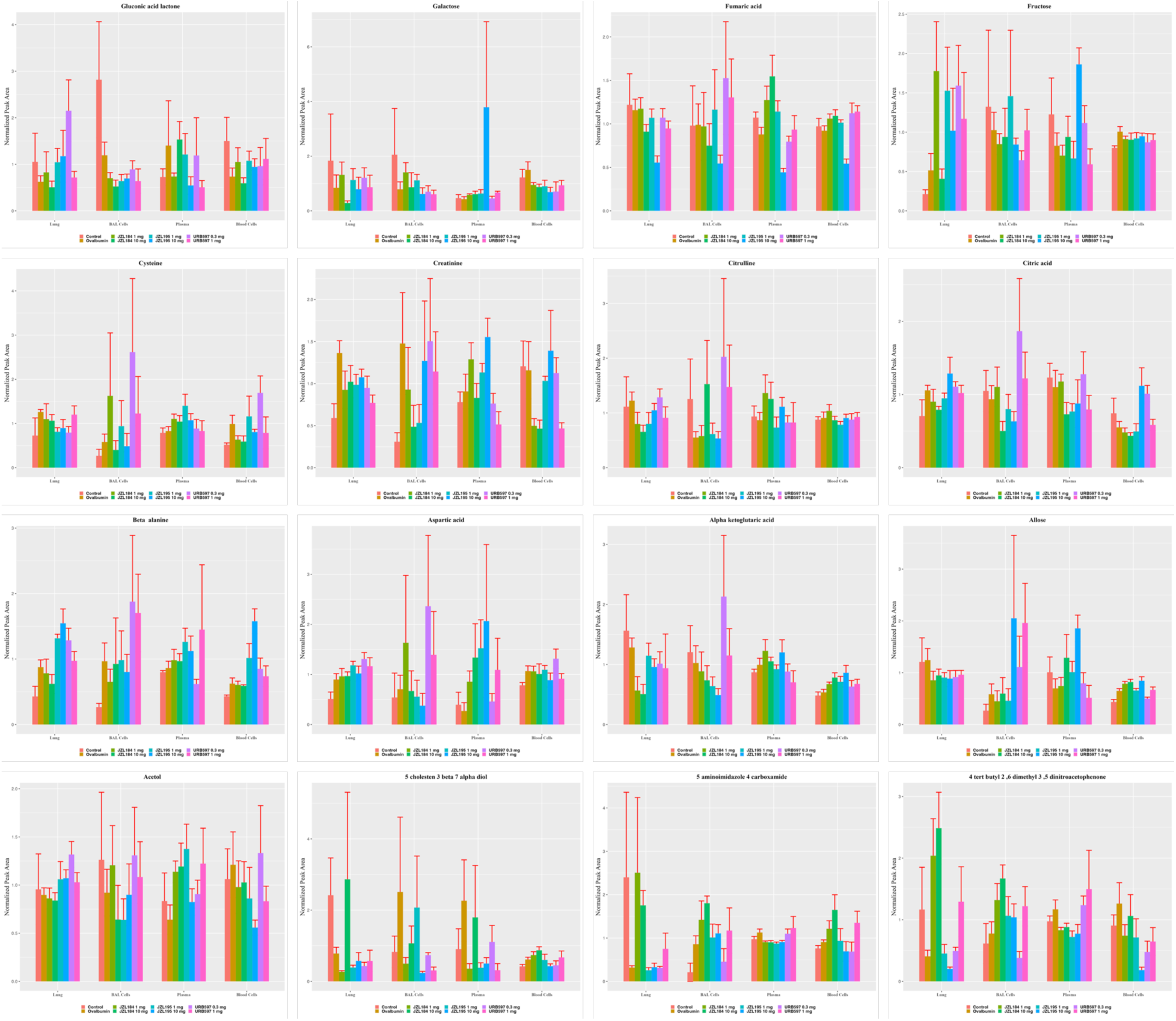
illustrates the alterations in different metabolites within the ovalbumin and treatment cohorts relative to the control group. The depicted changes are presented through bar graphs.

**Supplement Figure 11:**
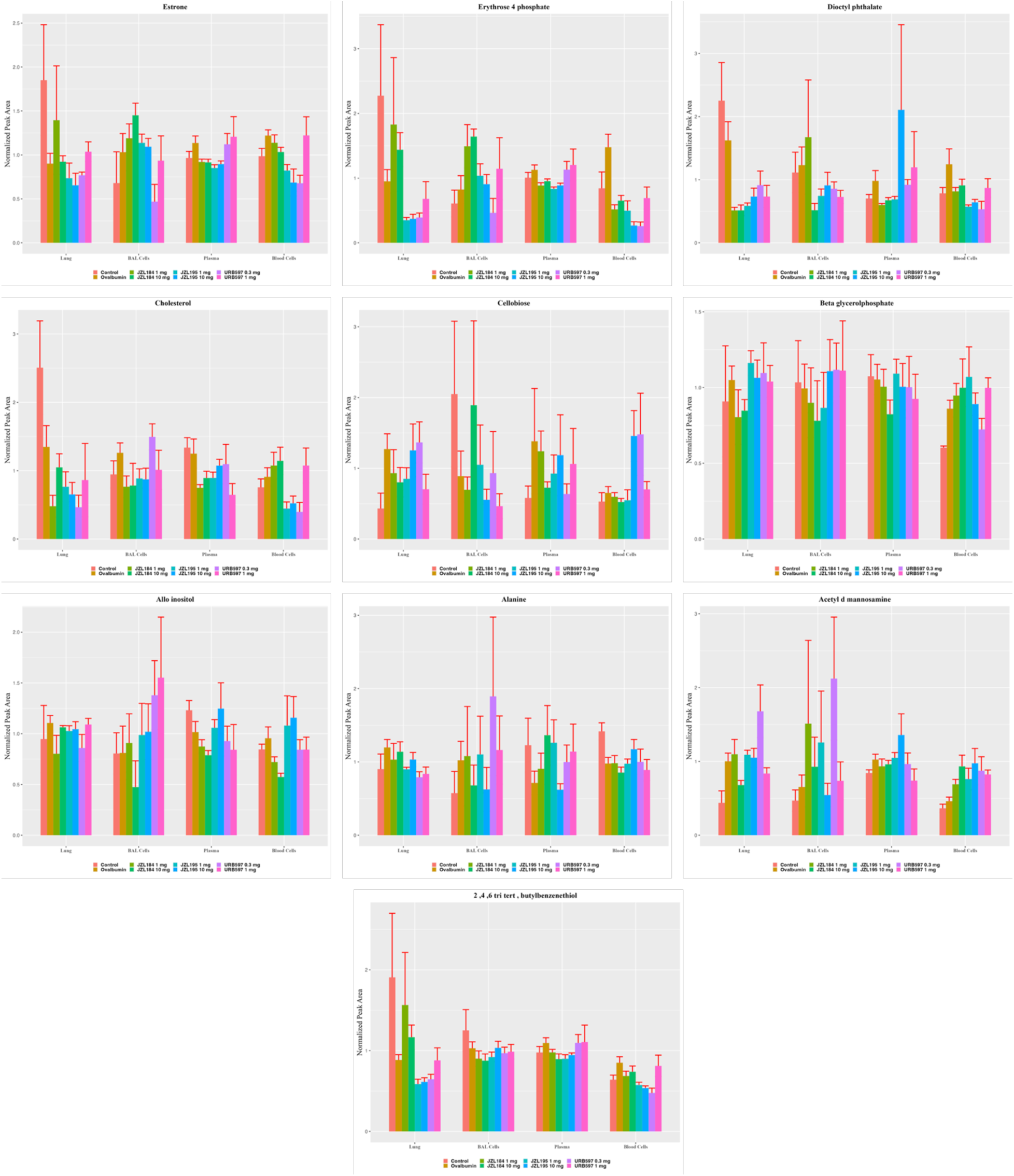
illustrates the alterations in different metabolites within the ovalbumin and treatment cohorts relative to the control group. The depicted changes are presented through bar graphs.

**Supplementary table 1:**
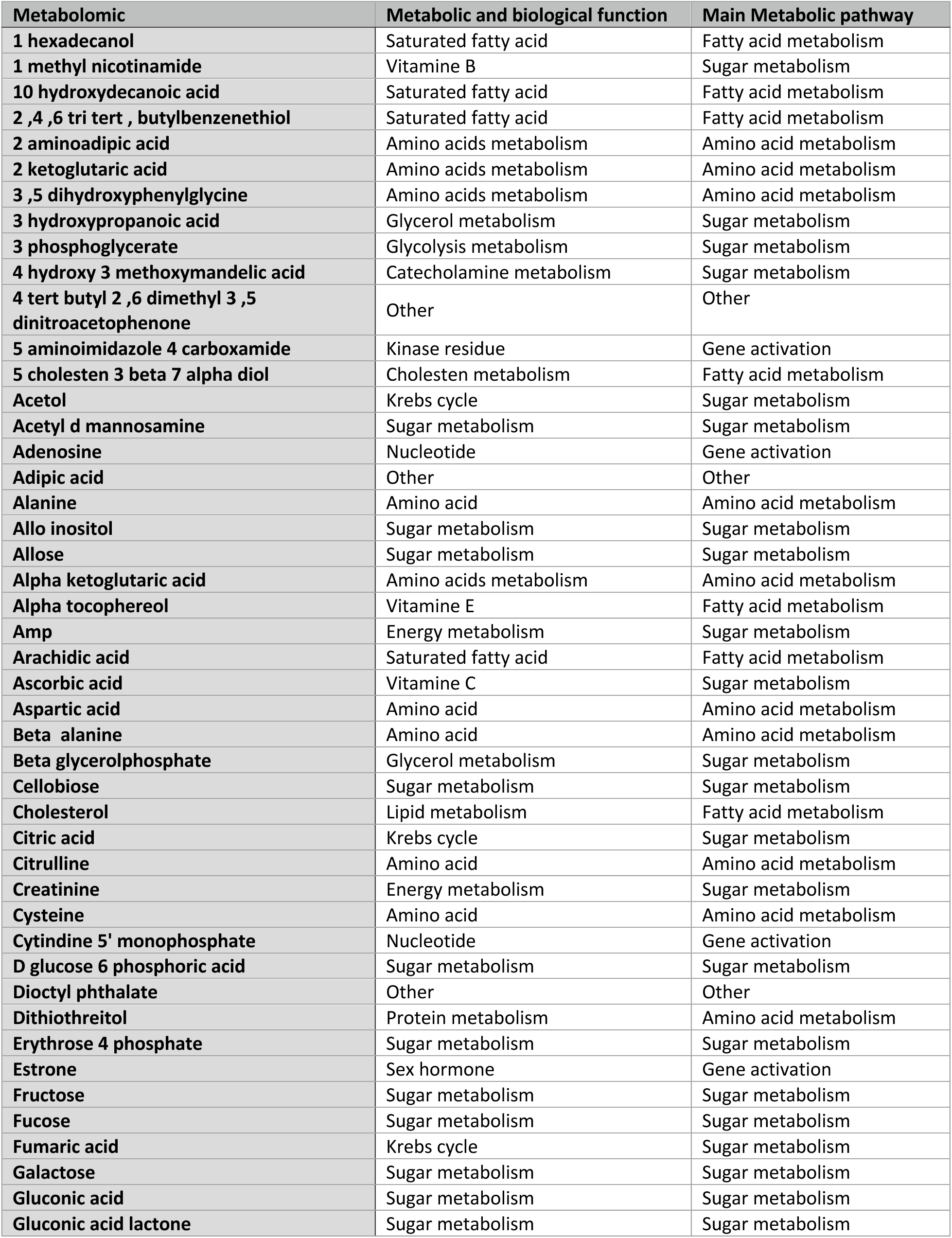
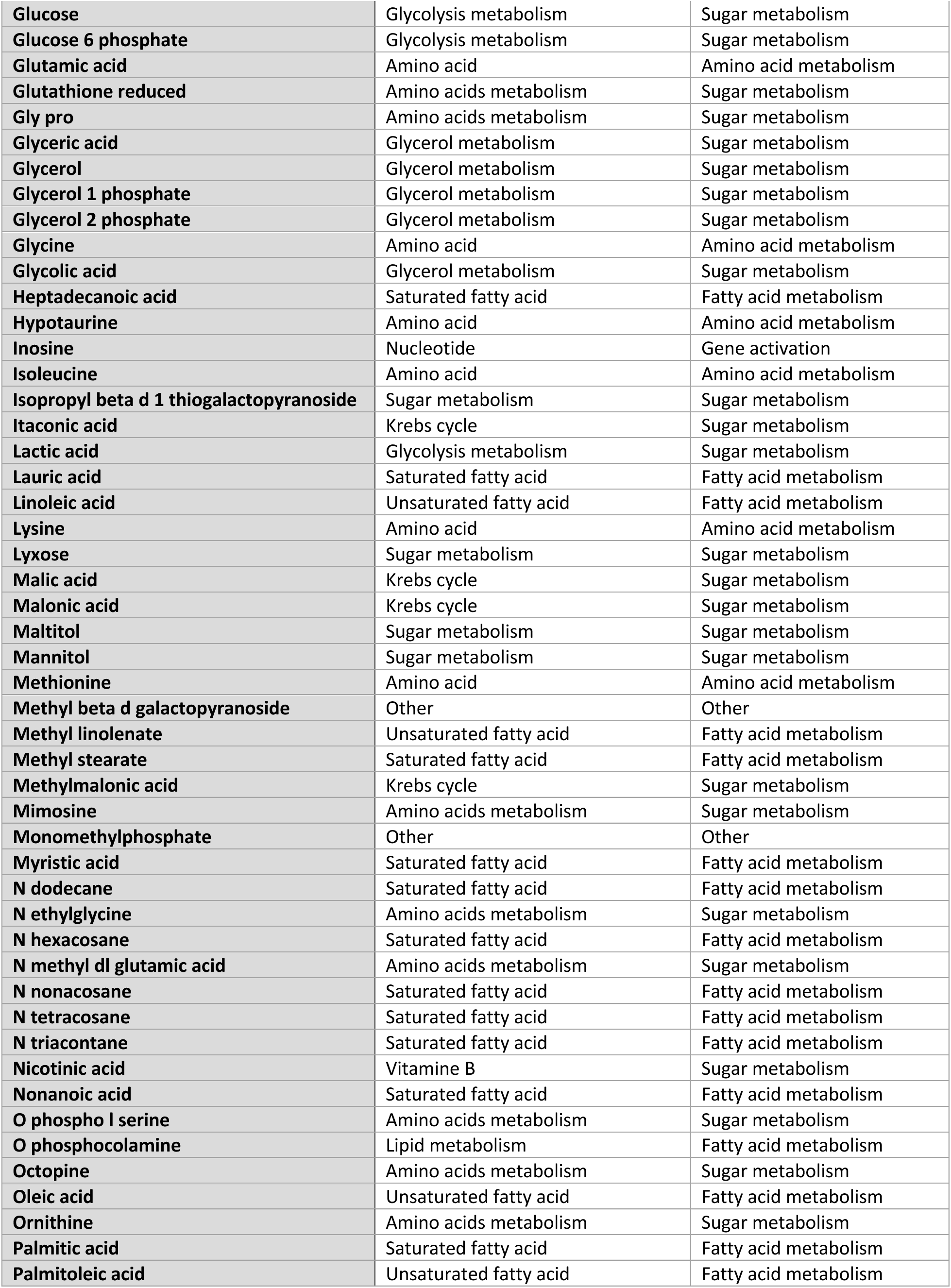
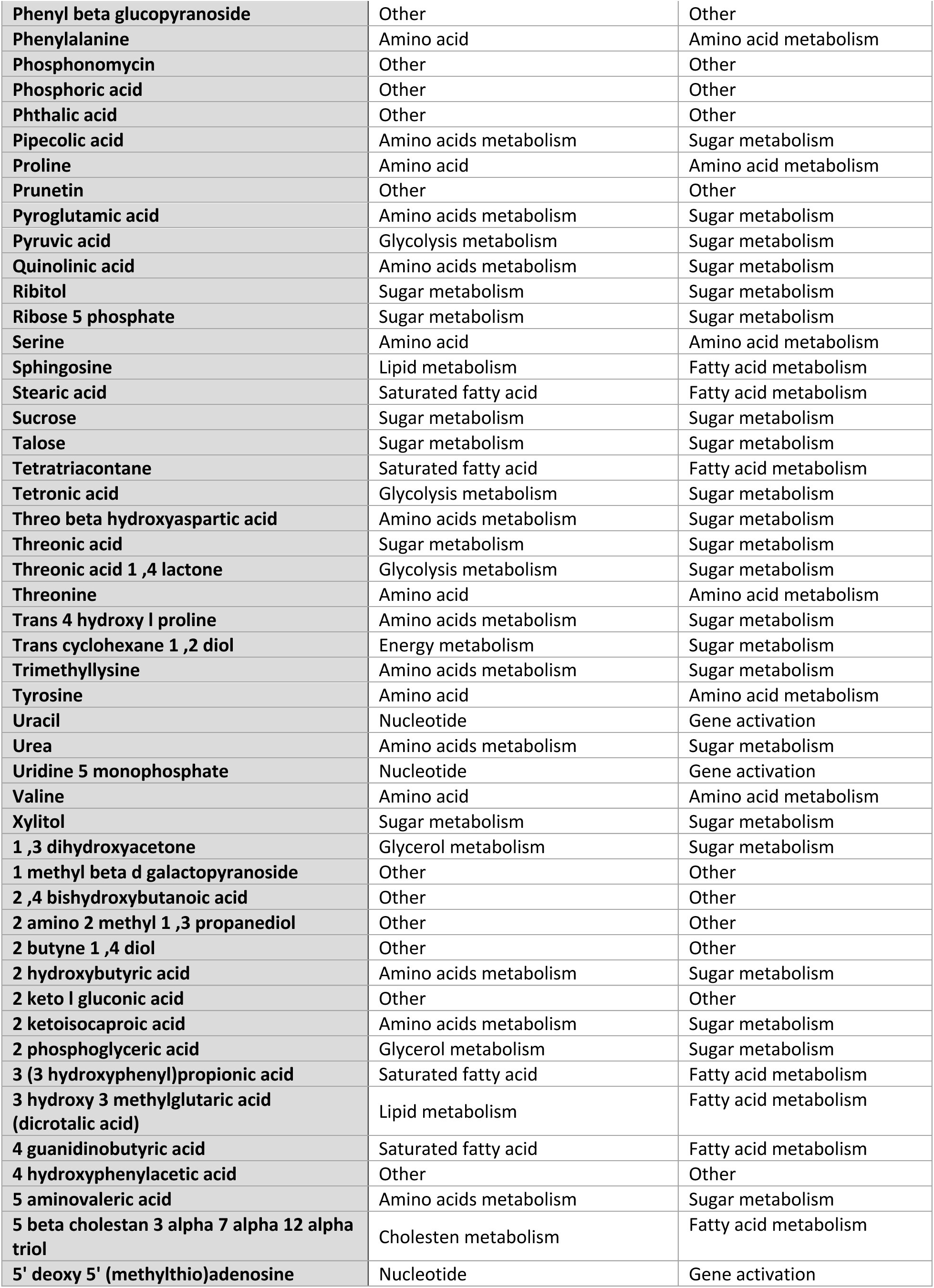
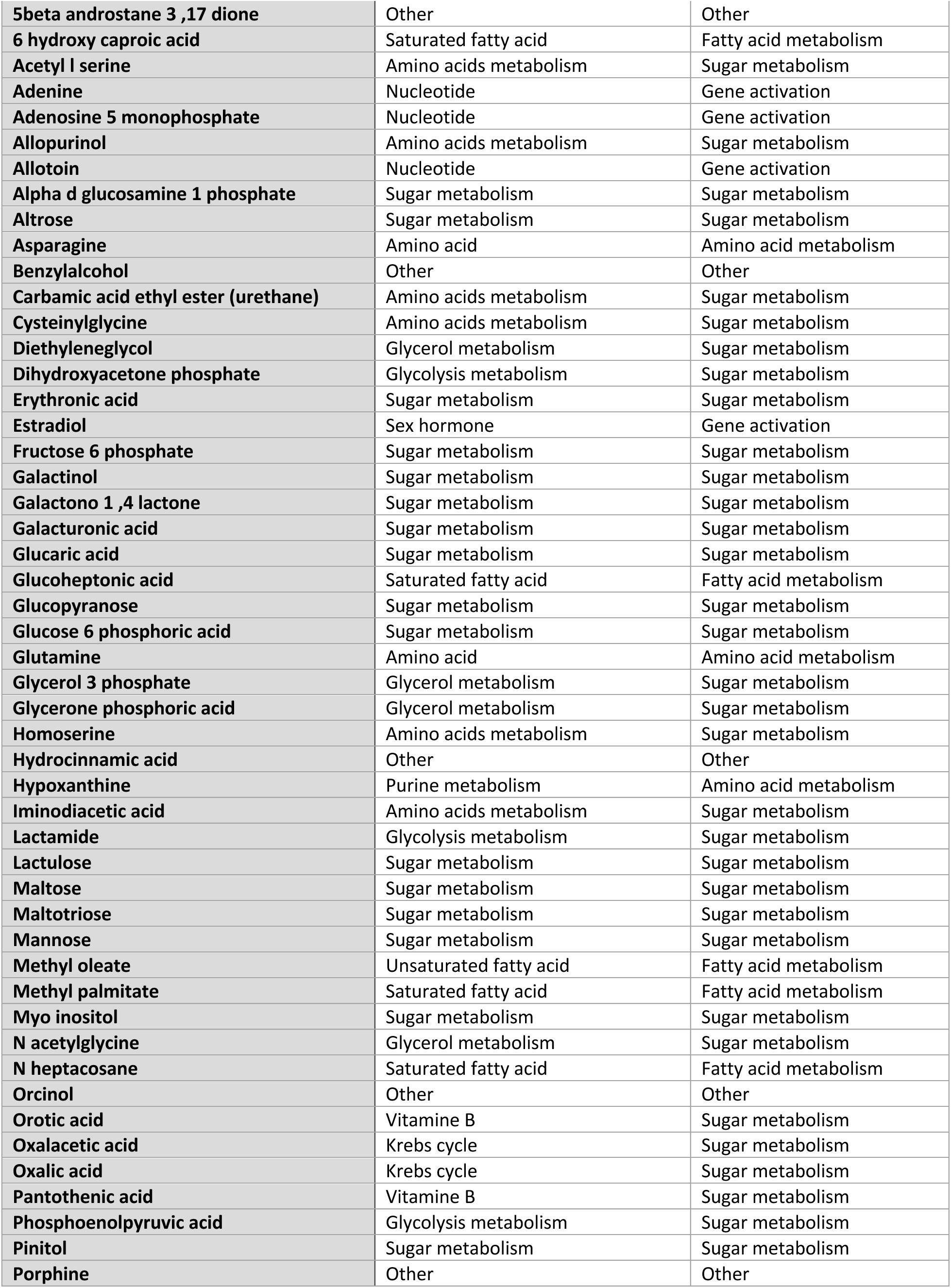
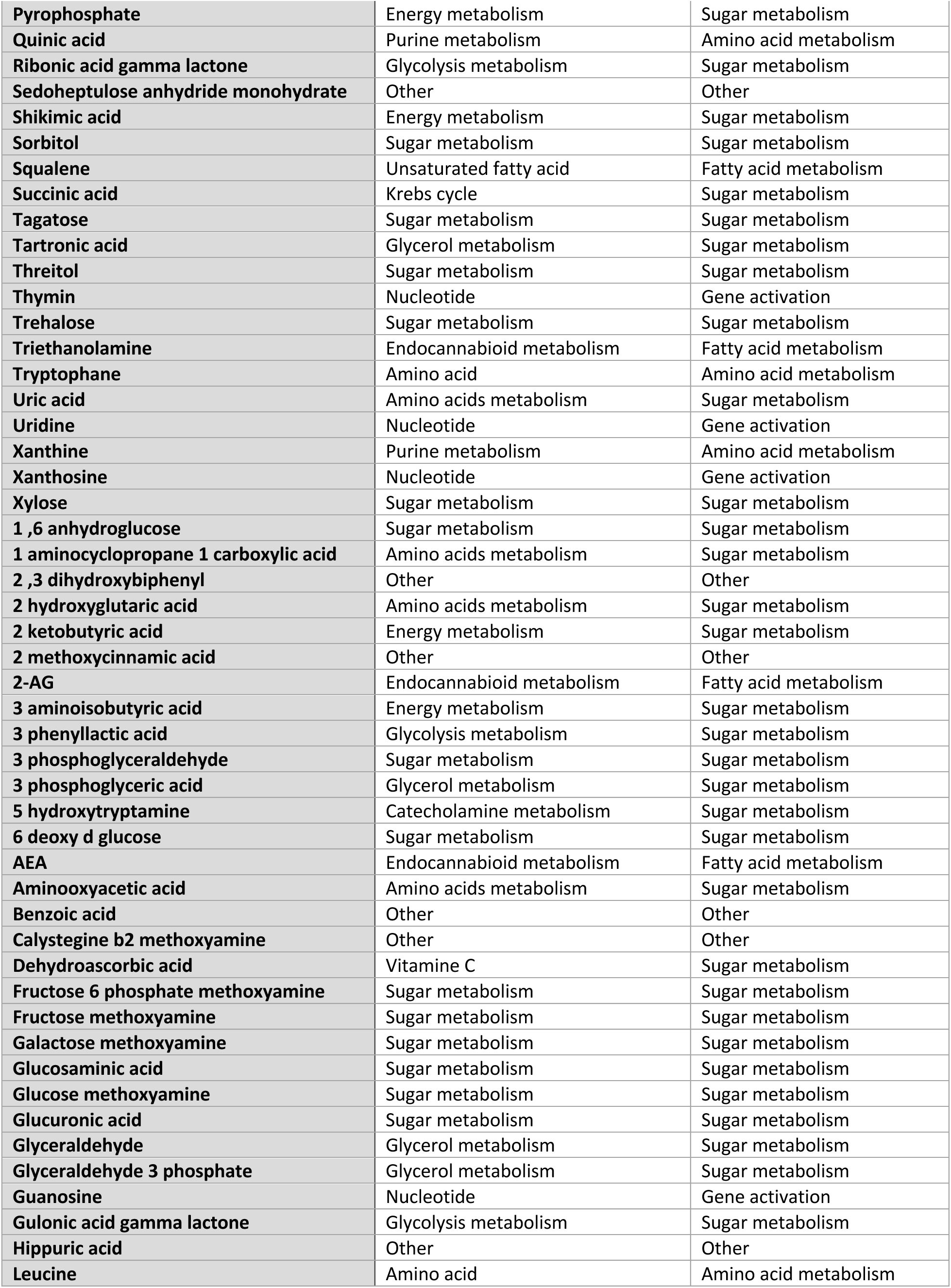
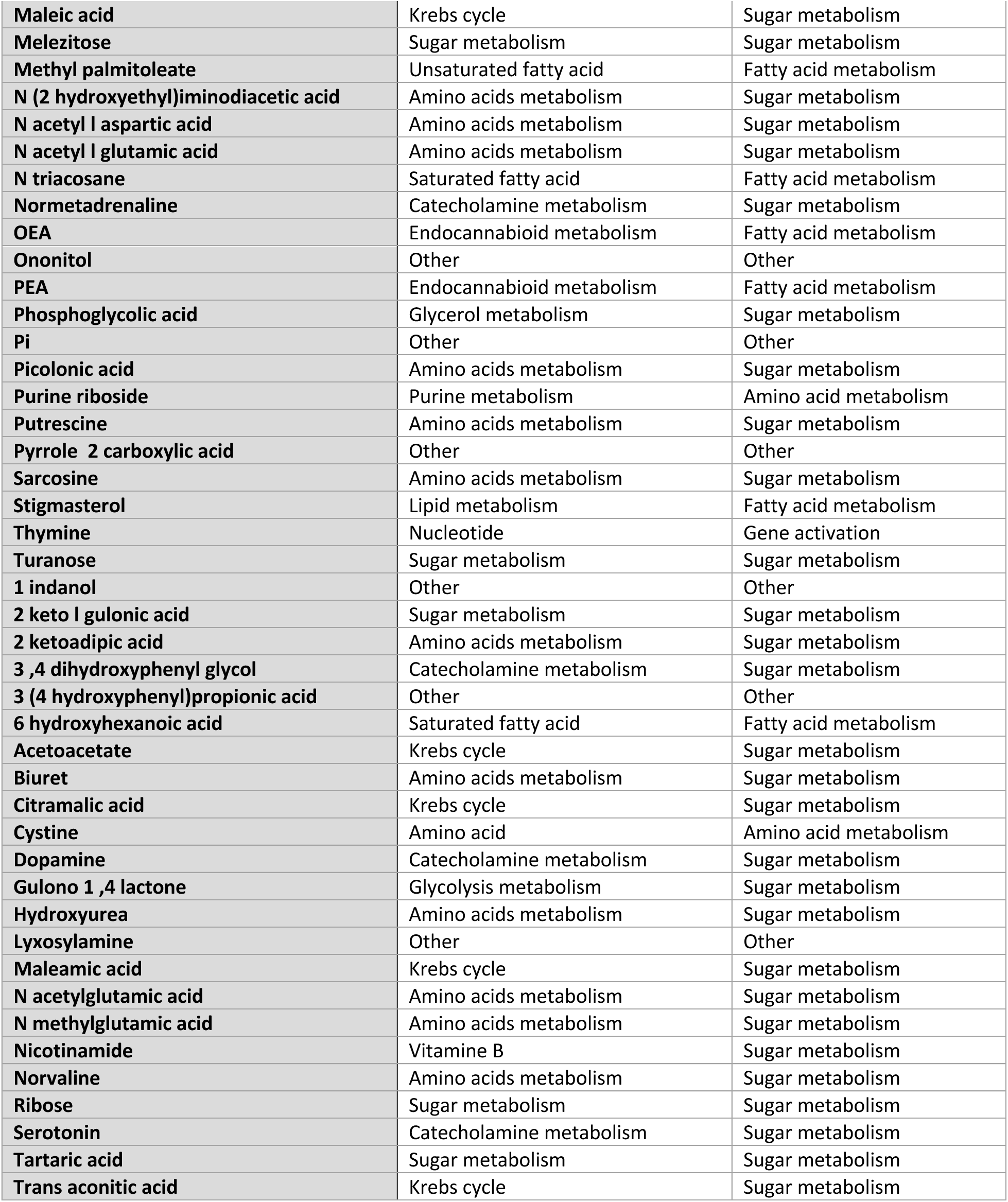
List of metabolomics classification.

## Notes

### Competing Interest Statement

The authors have declared no competing interest.

